# Neural Arbitration between Social and Individual Learning Systems

**DOI:** 10.1101/857862

**Authors:** Andreea O. Diaconescu, Madeline Stecy, Lars Kasper, Christopher J. Burke, Zoltan Nagy, Christoph Mathys, Philippe N. Tobler

**Affiliations:** Translational Neuromodeling Unit, Institute for Biomedical Engineering, University of Zurich & ETH Zurich, Switzerland; Laboratory for Social and Neural Systems Research, University of Zurich, Switzerland; University of Basel, Department of Psychiatry (UPK), Basel, Switzerland; Krembil Centre for Neuroinformatics, Centre for Addiction and Mental Health (CAMH), University of Toronto, Canada; Rutgers Robert Wood Johnson Medical School, New Jersey, United States; Institute for Biomedical Engineering, MRI Technology Group, ETH Zürich & University of Zurich, Switzerland; Scuola Internazionale Superiore di Studi Avanzati (SISSA), Trieste, Italy; Max Planck UCL Centre for Computational Psychiatry and Ageing Research, London, UK

**Author notes:** The authors contributed equally to this work (Andreea O. Diaconescu, Madeline Stecy). Correspondence should be addressed to: Andreea O. Diaconescu, PhD, Krembil Centre for Neuroinformatics (CAMH), 250 College St. M5T 1R8. **Disclosure statement:** The authors have no disclosures or conflict of interest.

**Keywords:** hierarchical Bayesian inference, observational learning, reinforcement learning, precision, uncertainty, fMRI, dopamine

## Abstract

Decision making often requires integrating self-gathered information with information acquired from observing others. Depending on the situation, it may be beneficial to rely more on one than the other source, taking into account that either information may be imprecise or deceiving. The process by which one source is selected over the other based on perceived reliability, here defined as arbitration, has not been fully elucidated. In this study, we formalised arbitration as the relative reliability (precision) of predictions afforded by each learning system using hierarchical Bayesian models. In a probabilistic learning task, participants predicted the outcome of a lottery using recommendations from a more informed advisor and self-sampled outcomes. The number of points participants wagered on their predictions reflected arbitration: The higher the relative precision of one learning system over the other and the lower the intention volatility, the more points participants wagered on a given trial. Functional neuroimaging demonstrated that the arbitration signal was independent of decision confidence and involved modality-specific brain regions. Arbitrating in favour of self-gathered information activated the dorsolateral prefrontal cortex and the midbrain whereas arbitrating in favour of social information engaged ventromedial prefrontal cortex and the temporoparietal junction. These findings are in line with domain specificity and indicate that relative precision captures arbitration between social and individual learning systems at both the behavioural and neural level.

## Introduction

As social primates navigating an uncertain world, humans use multiple information sources to guide their decisions (Charness et al., 2013). For example, in investment decisions someone may either choose to follow a financial expert’s advice about a particular stock or base the decision on their own previous experience with that stock. When information from personal experience and social advice conflict, one source must be favoured over the other to guide decision making. We conceptualize the process of selecting between information sources as arbitration. Arbitration is particularly important when the reliability of information is uncertain. While stock performance may fluctuate, the advisor could pursue selfish interests. It is challenging to infer the intentions of the advisor because they are concealed or expressed indirectly, requiring inference from observations of ambiguous behaviour. Optimal arbitration should therefore consider the relative uncertainty associated with each source of information.

Arbitration between different types of reward predictions based on experiential learning acquired by an individual has been associated with the prefrontal cortex. Specifically, the dorsolateral prefrontal cortex (DLPFC) and the frontopolar cortex have been shown to arbitrate between habitual (model-free) and planned (modelbased) learning systems (Lee et al., 2014). By contrast, comparatively little is known about how humans weigh self-gathered (individual) reward information against observed (social) information. To fill this gap, we considered two hypotheses: Arbitration involving social information could rely on theory of mind processes (ToM), i.e., inference about others’ mental states (Frith and Frith, 2005; Schaafsma et al., 2015) and higher-level social representations (Frith, 2012; Devaine et al., 2014a). Accordingly, arbitration involving the intentions of others may rely on activity in classical ToM regions, such as the temporoparietal junction (TPJ) and dorsomedial prefrontal cortex (Carrington and Bailey, 2009; Frith and Frith, 2010; Baker et al., 2011; Schurz et al., 2014). Alternatively, arbitration between individual and social information may involve similar neural networks as those orchestrating between model-free and model-based learning (Lee et al., 2014), and thus engage similar lateral prefrontal and frontopolar regions.

To investigate arbitration between individual and social learning systems, we simulated the aforementioned stock investment scenario in the laboratory. In other words, we examined how people arbitrate between individual reward information and social advice regarding a probabilistic lottery where contingencies changed over time. Participants learned to predict the colour of a binary card draw using uncertain advice from a more informed advisor and uncertain information inferred from individually observed card outcomes (Figure 1). We separately manipulated the degree of uncertainty (or its inverse, precision) associated with each of the two information sources by independently varying the rate of change with which each information source predicted the drawn card colour (i.e., volatility; Behrens et al., 2007). The advisor was motivated to give correct or incorrect advice depending on the phase of the task, resulting in variations in the reliability of social information. Performing well in the task therefore required keeping track of the probabilities of the two sources of information and arbitrating between them. We assumed that participants weigh the predictions afforded by each information source as a function of their precision. Thus, we expected participants to rely more on the advice when the advisor’s intentions were perceived as stable, and on their personal experience when the advice was perceived to be volatile.

**Figure 1.**
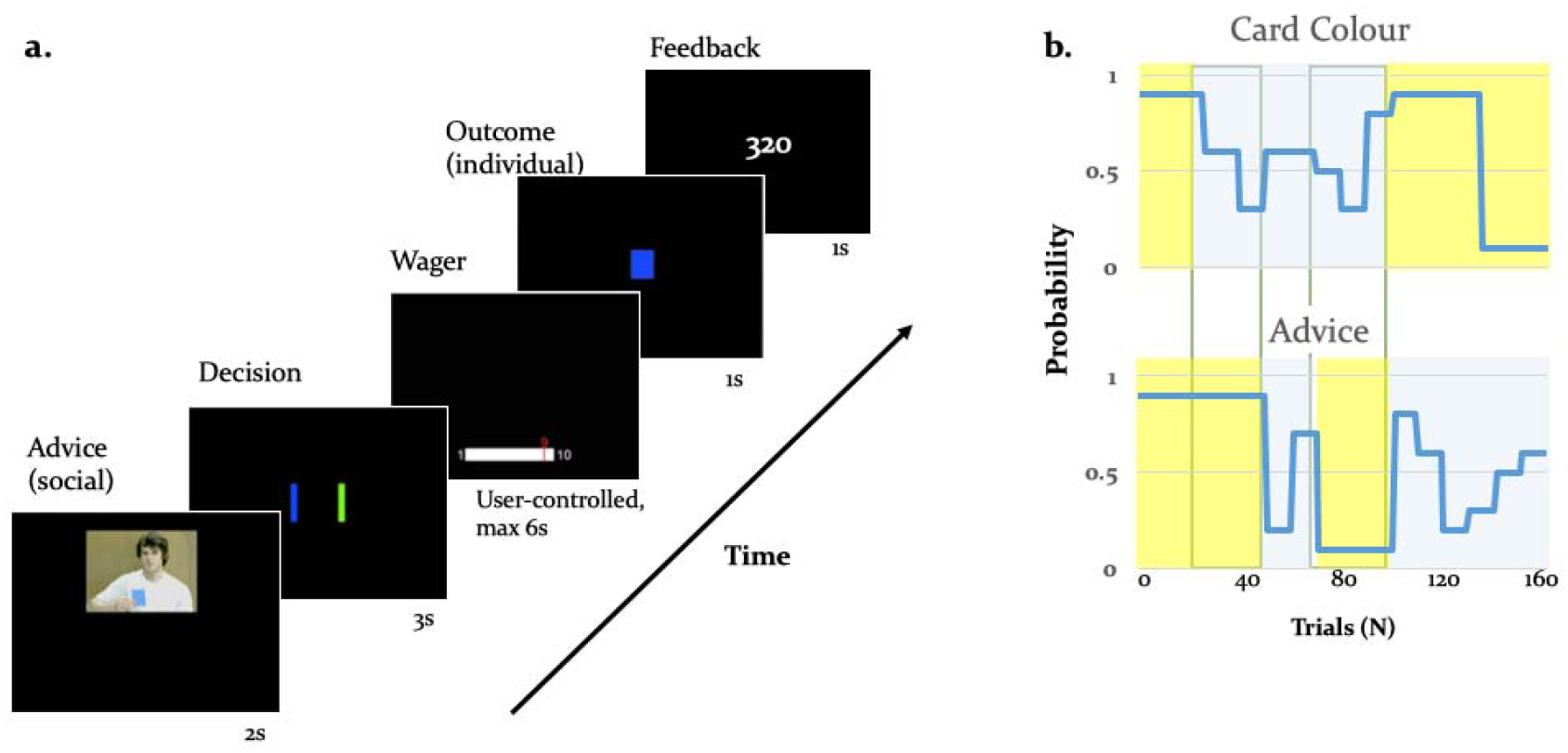
Experimental Paradigm: (a) Binary lottery game requiring arbitration between individual experience and social information. Volunteers predicted the outcome of a binary lottery, i.e., whether a blue or green card would be drawn. They could base the prediction on two sources of information: advice from a gender-matched advisor (video, presented for 2s) who was better informed about the colour of the drawn card, and on an estimate about the statistical likelihood of the cards being one or the other colour that the subject had to infer from own experience (outcome, 1s). After predicting the colour of the rewarded lottery card (user-controlled, maximum 3s), subjects also wagered one to ten points (user-controlled, maximum 6s), which they would win or lose depending on whether the prediction was right or wrong. After the outcome participants viewed their cumulative score on the feedback screen (1s), (b) Contingencies of individual reward and social advice information. Card colour probability corresponds to the likelihood of a given colour (e.g., blue) being rewarded. The two sources of information were uncorrelated as illustrated by phases of low (yellow) and high (blue) volatility, enabling a factorial analysis of information source and volatility.

## Results

To examine the neural mechanisms underlying arbitration, we combined fMRI with a computational modelling approach using the hierarchical Gaussian filter (HGF) (Mathys et al., 2011, 2014). This hierarchical Bayesian model is ideally suited to address our question as it examines multi-level inference and provides trial-wise estimates of estimated precision of predictions about each information source. This framework operationalises arbitration as a precision ratio, corresponding to the relative perceived precision of each information source (Figure 2). Thus, arbitration is a function of the relative stability of the advice or the card colour probabilities. In our paradigm, this quantity increased when the precision of the predictions about one of the two sources of information was high and decreased when both sources were either stable or volatile (see Figure 4c for the arbitration signal averaged across participants).

**Figure 2.**
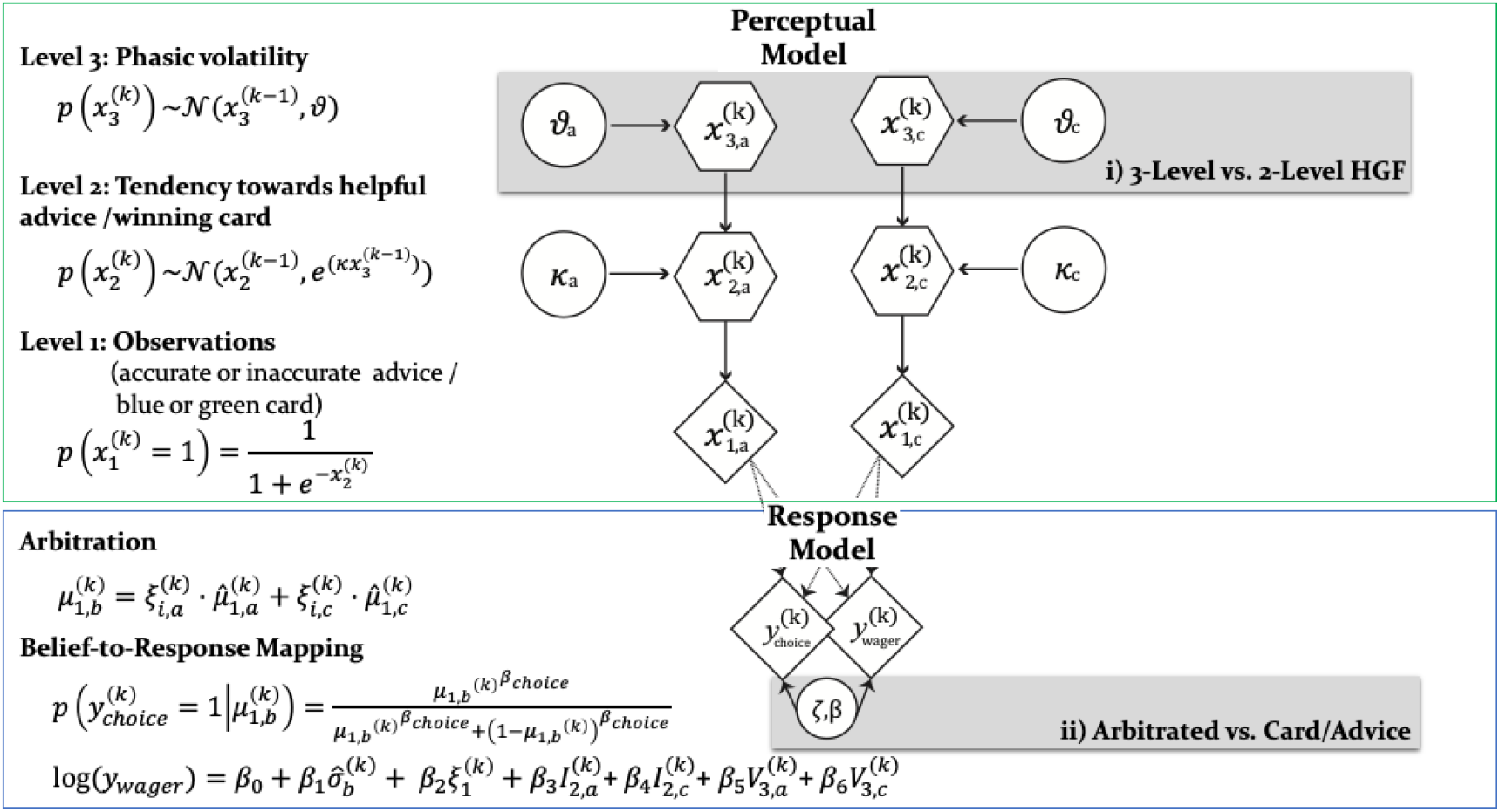
Computational learning and arbitration model: In this graphical notation, circles represent constants whereas hexagons and diamonds represent quantities that change in time (i.e., that carry a time/trial index). Hexagons in contrast to diamonds additionally depend on the previous state in time in a Markovian fashion. Two parallel HGFs describe the generative model for advice and card probability: x_1_ represents the accuracy of the current advice / card colour probability, x_2_ the tendency of the advisor to offer helpful advice / tendency of card colour to be rewarded, and x_3_ the current volatility of the advisor’s intentions / card colour probabilities. Learning parameters describe how the states evolve in time. Parameter κ determines how strongly x_2_ and x_3_ are coupled, and □ represents the metavolatility of x_3_. The response model maps the predicted colour probabilities to choices. For model selection we combined two perception with three response models (see Figure 3). All the models considered can be grouped according to common features and divided into model families: (i) the Perceptual model families distinguish between hierarchical (3-level) and non-hierarchical (2-level) HGFs, which refer to estimating or fixing the volatility of the third level. The 3-level HGF model assumes that trial-wise wagers and predictions arise from a linear combination of arbitration, informational uncertainty (advice and card), and volatility (advice and card), (ii) Response model families distinguish between arbitrated and single-information source – advice or card only – models, which correspond to estimating parameter *ζ* or fixing it to reduce arbitration to either the advice prediction or the card colour prediction.

### Behaviour: Prediction accuracy and wager size

Using the factorial structure of the task, we tested the impact of volatility on performance with a two-factor repeated measures ANOVA, where the two factors were information source (card versus advice) and phase (stable versus volatile). Across all behavioural metrics, we observed an effect of phase, indicating a reduction in performance in volatile compared to stable phases, and a phase × cue interaction, indicating that the effect was larger for the social than the individual source of information. First, for prediction accuracy, we found a main effect of phase (*df* = (1,36), *F* = 187.94, *p* = 7.7e-16) and an information source-by-phase interaction (*df* = (1,36), *F* = 11.13, *p* = 0.0020) (see Figure S1a). Thus, in-keeping with the reasoning that arbitration relates to relative information quality, the degree to which participants relied on each information source was a function of precision as manipulated by the volatility structure of the task. Participants performed significantly better in stable compared to volatile periods of the task. These effects were not modulated by fatigue, as we found no significant differences between early and late phases of the task.

**Figure S1.**
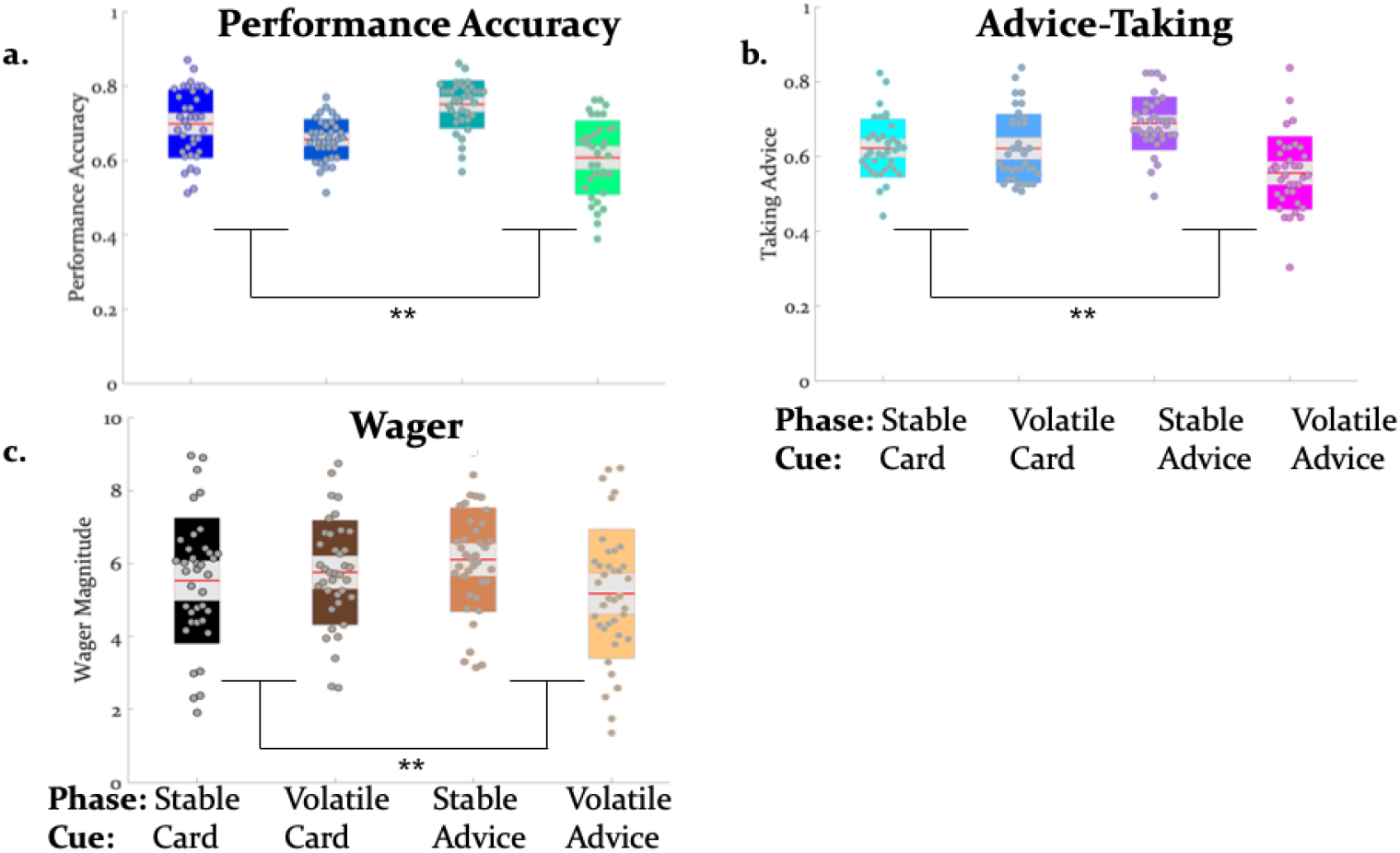
Behaviour influenced by volatility: Average prediction accuracy (a), advice-taking behavior (b), and amount of points wagered per trial (c) were reduced during volatile phases of the paradigm, particularly with regard to social information. The average values across all trials were 68.2%± 6.2% (mean accuracy ± standard deviation) prediction accuracy, 62.1%± 6.9% advice-taking, and 5.6 ± 1.5 points wagered (participants on average accumulated 378.6± 173.2 points). Jittered raw data (i.e., means over all trials of each behavioural measure per subject) are plotted for each behavioural measure. Red lines indicate the mean, grey areas reflect 1 SD of the mean, and coloured areas the 95% confidence intervals of the mean. \*\**p* <0.001 is indicated to emphasize the phase × cue interactions.

Second, also advice-taking behaviour differed as a function of volatility and information source: For the percentage of trials in which participants followed a given source of information, we detected a main effect of phase (*df* = (1,36), *F* = 56.26, *p* = 7.3073e-09) and an information source-by-phase interaction (*df* = (1,36), *F* = 25.86, *p* = 1.1561e-05) (Figure S1b). Thus, participants took advice less often particularly when it was volatile rather than stable.

Third, the amount of points wagered also depended on volatility and information source. We observed a main effect of phase (*df* = (1,36), *F* = 28.78, *p* = 4.54e-06) and an information source-by-phase interaction (*df* = (1,36), *F* = 16.75, *p* = 2.21e-04 (Figure S1c). Participants wagered fewer points particularly when advice was volatile. Moreover, the number of points wagered correlated significantly with the total score in stable phases (*r* = 0.37, *p* = 0.02), but not in volatile phases (*r* = 0.30, *p* = 0.06). Simulations using a nonvolatility, 2-level HGF suggested that tracking volatility is beneficial for task performance: a hypothetical person who did not take the volatility of the task phases into account gained on average 21.6 less points than an agent tracking volatility. In line with previous evidence (Behrens et al., 2008), these results emphasize the impact of volatility on the willingness to invest and investment success as measured here by total score.

### Model-based results

#### Model Selection

We used computational modelling with hierarchical Gaussian Filters (HGF; Figure 2) to explain participants’ responses on every trial. To contrast competing mechanisms underlying learning and arbitration, our model space included a total of 6 models (Figure 3a), with perceptual models varying in complexity of volatility processing (3-level full HGF vs. 2-level no-volatility HGF) and response models varying in the extent of arbitration (arbitration; no arbitration: advice only; no arbitration: card information only). Bayesian model selection (Stephan et al., 2009) served to compare models (see Methods and Figure 2 for details).

**Figure 3.**
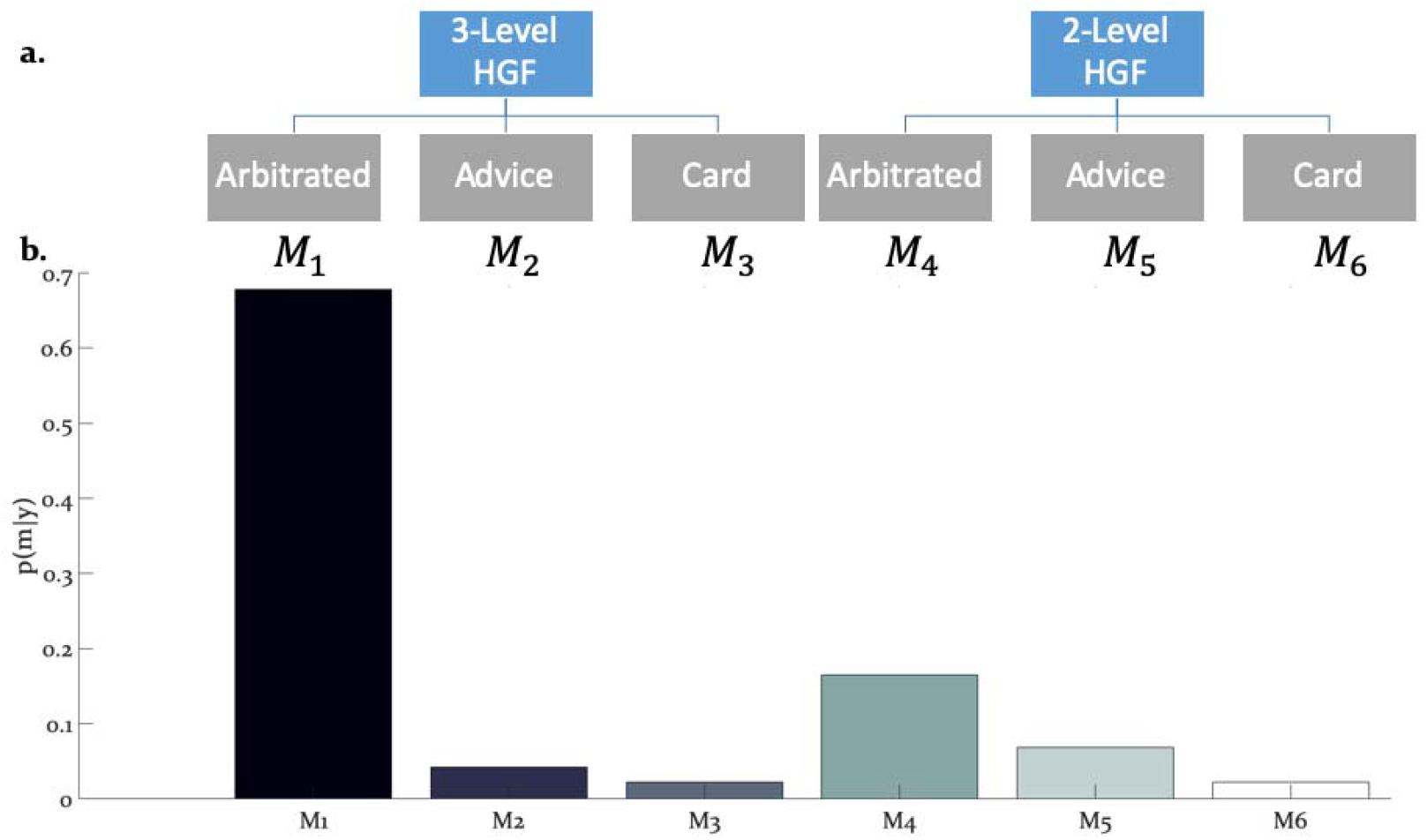
Hierarchical structure of the model space and model selection results: (a) The learning and arbitration models considered in this study have a 2 × 3 factorial structure and can be displayed as a tree. The nodes at the top level represent the perceptual model families (3-level HGF, 2-level nonvolatility HGF). The leaves at the bottom represent response models which integrate and arbitrate between social and individual sources of information (“Arbitrated”) or exclusively consider social (“Advice”) or individual (“Card”) information. (b) Bayesian model selection revealed one winning model, the Arbitrated 3-level HGF. Posterior model probabilities or *p*(*m*|*y*) revealed that this model explained participants’ behaviour in the majority of the cases.

The winning model was the 3-level HGF with arbitration (*ϕ_p_* = 0.999; Bayes Omnibus Risk = 2.0863e-10; Figure 3b; Table 2a). This model captured arbitration with the ratio of precisions: the precision of the prediction about advice accuracy and colour probability, divided by total precision. Moreover, the model included a social bias parameter reflecting the degree to which participants followed the advisor irrespective of task information. The model family that included volatility of both information sources outperformed models without volatility, in-keeping with the model-independent finding that perceived volatility of both information sources affected behaviour.

#### Posterior Parameter Estimates

Three parameters modulated the arbitration signal of the winning model. These included: (i) *κ* or the coupling between the two hierarchical levels that determined the impact of volatility on the inferred predictions of each information source (Eq. 6), (ii) *ϑ*, determining the variance of the volatility (Eq. 12), and (iii) *ζ*, the social bias which reflected the reliance on the advice independent of its reliability (Eq. 19). Both coupling *κ* and volatility parameter *ϑ* did not significantly differ between learning from individual and social information (t(36)= 0.53, p=0.60 and t(36)= −0.32, p=0.75 for *κ* and *ϑ*, respectively; Figure 4a). In fact, they were highly correlated: r_1_=0.55, p_1_=0.003 for *κ* and r_2_=0.64, p_2_=0.001 for *ϑ*. This result suggests that participants learned from individual (volatile card probabilities) and social (advisor fidelity) information similarly.

**Figure 4.**
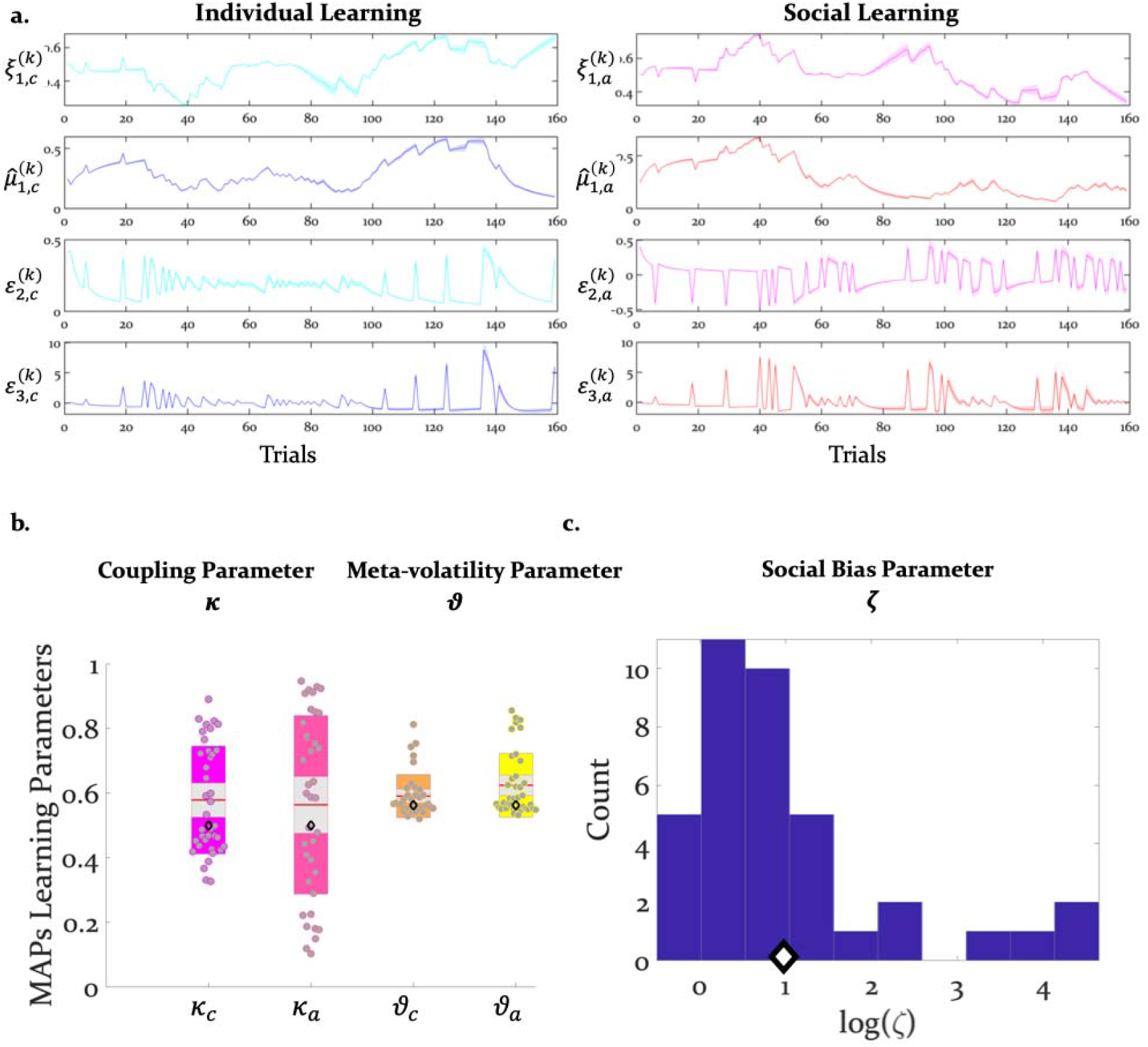
Inference and arbitration of individual and social learning: (a) Average trajectories for arbitration and hierarchical precision-weighted PEs for individual and social learning (see Methods for the exact equations): *ξ_a_* = arbitration in favour of the advice (Eq. 19); *ξ_c_* = arbitration in favour of individually estimated card colour probability (Eq. 20). 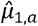 = estimated advice accuracy (Eq. 4); 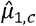 = individually estimated card colour probability (Eq. 18). *ε*_2,*a*_ = precision-weighted prediction error (PE) of advisor fidelity (Eq. 8); *ε*_2,*c*_ = unsigned (absolute) precision-weighted PE of card outcome (absolute value of Eq. 14). *ε*_3,*a*_ = precision-weighted advice volatility PE (Eq. 13); *ε*_3,*c*_ = precision-weighted card colour volatility PE (Eq. 15). Line plots are generated by averaging computational trajectories extracted from the winning (Arbitrated 3-HGF: Figure 2) model for each subject across trials, i.e. 0 to 160 trials. The shaded area around each line depicts +/- standard error of the mean over subjects. (b) Group means, standard deviations and prior values for the perceptual model parameters determining dynamics of computational trajectories in (a). (c) the distribution of log(zeta) values along with its prior. Jittered subject-specific estimates are plotted for each perceptual model parameter, red lines indicate the group mean, grey areas reflect 1 SD of the mean, and coloured areas the 95% confidence intervals of the mean. In (b) and (c), black diamonds denote the priors of each parameter (for details, see Table 1).

The reliability-independent social bias parameter ζ did not differ significantly from an unbiased value of 1 (t(36)= −0.14, p=0.89). Thus, on average, we found no dominant strategy of preferentially selecting or dismissing the advisor’s suggestion (Figure 4b). Still, participants varied in how much they relied on advice to predict card colour (Figure 4c).

According to the model, decisions of how many points to wager on a given trial related to several time-varying factors. These included the (irreducible) uncertainty of the agent’s beliefs about the decision, arbitration and the estimated volatility of the advisor’s intentions (belief uncertainty: t(37)=6.64, p=2e-12; arbitration: t(37)=6.64, p=6e-06; and estimated advisor volatility: t(37)=−7.11 p= 2e-08) (Figure 5). The higher the decision confidence and the stronger the bias to arbitrate in favour of social information, the more points participants wagered. Conversely, estimated advisor volatility was negatively associated with the amount wagered: the higher the estimated advisor volatility, the fewer points participants were willing to wager on a given trial (see Table 1 for the priors over the parameters, Table 2b for all parameter estimates, and Figure 5 for the trial-wise influence of the average computational quantities on wager amount).

**Figure 5.**
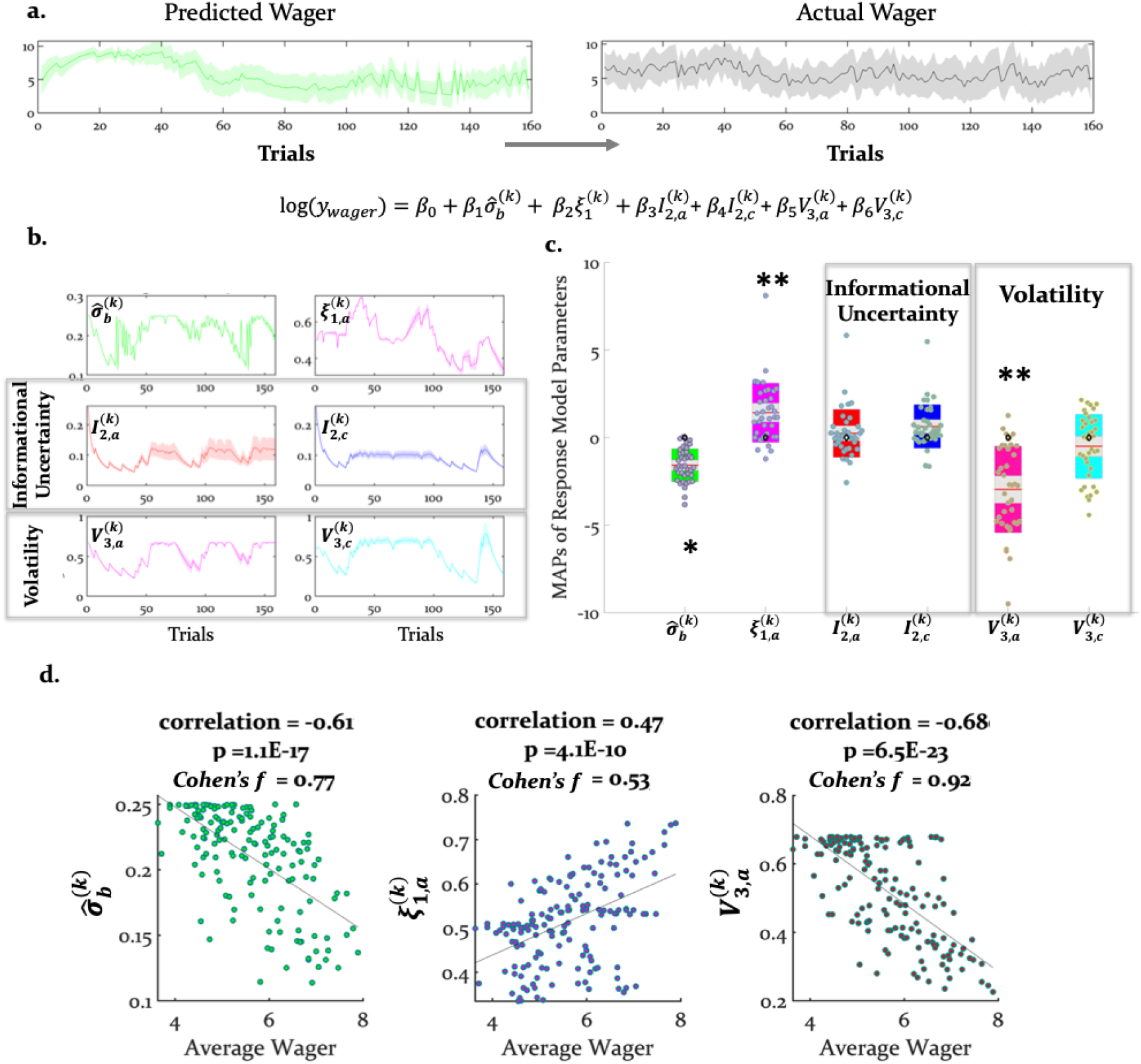
Predicting the wager: Computational quantities and model parameters influencing wagers: (a) With our response model, we predicted that the actual trialwise wager (right) could be explained (left and bottom) by 6 key trajectories (see Eqn. 21). (b) These include (i) (irreducible) belief uncertainty or 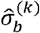 (based on the integrated belief of individual and advice predictions; Eqn. 24); (ii) arbitration in favour of advice (Eqn. 19); (iii) informational uncertainty (Eqn. 24) and volatility of the advice (Eqn. 25) and (iv) informational uncertainty (Eqn. 26) and volatility of the card (Eqn. 28) in (b). (a) and (b) show group averages (see Methods for the exact equations). For the model-based parameters, the line plots are generated by averaging computational trajectories extracted from the winning model (Arbitrated 3-HGF) for each subject across trials, i.e. 0 to 160 trials. The shaded areas depict +/- standard error of this mean over subjects. (c) Group means, standard deviations and prior values for the response model parameters determining the impact of those trajectories (i.e., uncertainties and arbitration) on trial-wise wager amount. Jittered raw data are plotted for each parameter. Red lines indicate the mean, grey areas reflect 1 SD of the mean, and the coloured areas the 95% confidence intervals of the mean. The black diamonds denote the prior of the parameters, which in this case is zero. \**p*<0.05, \*\**p*<0.001. (d) Scatter plots with average actual wager on the x-axis and average belief uncertainty, arbitration in favour of advice, and volatility of advice on the y-axes, respectively. The correlation coefficients (with corresponding p values), regression slopes, and effect sizes (Cohen’s *f*) are included to quantify the relationship between the actual wager and the computational quantities we found to significantly predict wagers.

**Table 1:** Prior mean and variance of the perceptual and response model parameters

| Model |  | Prior mean | Prior variance |
| --- | --- | --- | --- |
| **Perceptual Models:** |  |  |  |
| | $\kappa_a, \kappa_c$ | 0.5 | 1 |
| **3-level HGF** | $\vartheta_a, \vartheta_c$ | 0.62 | 1 |
| **2-level HGF** | $\vartheta_a, \vartheta_c$ | 0.00062 | 0 |
| **Response Models:** |  |  |  |
| | $\beta_{1-6}$ | 0 | 4 |
| | $\beta_{ch}$ | 48 | 1 |
| | $\beta_0$ | 6.21 | 4 |
| | $\beta_{wager}$ | 1.50 | 100 |
| **1. Arbitrated** | $\zeta$ | 1 | 25 |
| **2. Advice Only** | $\zeta$ | Inf | 0 |
| **3. Card Only** | $\zeta$ | 0 | 0 |

Note: The prior variances are given in the numeric space in which parameters are estimated. $\kappa$ , $\vartheta$ , and $\mu_3^{(k=0)}$ are estimated in logit-space, while the other parameters are estimated in log-space. Whereas the prior variances for all parameters are set to be rather broad, we selected a shrinkage prior mean and variance for the decision noise parameter $\beta_{ch}$ such that behaviour is explained more by variations in the remaining parameters rather than decision noise.

**Table 2a:** Results of Bayesian model selection: Model probability (***p***(***m***|***y***)) and protected exceedance probabilities (***ϕ_p_***)

| Response Models: | *Perceptual Models:*<br>*3-level HGF* |  |  |
| --- | --- | --- | --- |
|  | Arbitrated | Advice Only | Card Only |
| $p(m\vert y)$ | 0.68 | 0.04 | 0.02 |
| $\phi_p$ | 0.99 | 1.0215e-10 | 1.0215e-10 |
|  | *2-level HGF* |  |  |
| $p(m\vert y)$ | 0.17 | 0.04 | 0.02 |
| $\phi_p$ | 5.9000e-05 | 1.0215e-10 | 1.0215e-10 |

**Table 2b:** Average maximum a-posteriori estimates of the learning and arbitration parameters of the winning model (Arbitrated 3-level HGF)

| Perceptual Model Parameters | Mean | SD | Response Model Parameters | Mean | SD |
| --- | --- | --- | --- | --- | --- |
| $\kappa_c$ | 0.61 | 0.17 | $\zeta$ | 0.98 | 1.17 |
| $\vartheta_c$ | 0.64 | 0.23 | $\beta \xi$ | 1.60 | 1.78 |
| $\kappa_a$ | 0.59 | 0.27 | $\beta \hat{\sigma}_1$ | -1.62 | 0.94 |
| $\vartheta_a$ | 0.66 | 0.27 | $\beta \hat{\sigma}_{2,a}$ | 0.13 | 1.45 |
| | | | $\beta \hat{\sigma}_{2,r}$ | 0.59 | 1.17 |
| | | | $\beta \hat{\mu}_{3,a}$ | -2.98 | 2.51 |
| | | | $\beta \hat{\mu}_{3,r}$ | -0.65 | 1.82 |
| | | | $\beta_{ch}$ | 2.27 | 0.92 |

#### Internal Validity of the Model

We aimed to examine at the behavioural level whether the model predictions were consistent with participants’ perceptions of the advice accuracy during the experiment. Participants judged advice accuracy (i.e., helpful, misleading, or neutral with regard to predicting actual card colour) in a multiple-choice question presented 5 times during the experiment (initial/prior: 1^st^ trial; stable advice, stable card phase = (14^th^ trial); stable advice, volatile card phase (49^th^ trial); volatile advice, volatile card phase (73^rd^ trial); volatile advice, stable card phase = 115^th^ trial). We first tested whether the responses to these questions positively related to estimates of advice accuracy 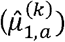 that were extracted from the winning model. A linear regression analysis demonstrated that the inferred advice accuracy or 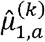 measured at the time of the multiple-choice question, predicted participants’ selections. Specifically, the estimated beta parameter estimate across all task phases was significantly different from zero (t(36) = 4.71, p = 3e-05). These findings suggest that the model predicted independently (and discretely) measured perception of advice accuracy, in-keeping with the internal validity of the model.

Next, we tested whether the wager magnitudes predicted by the model correlated with participants’ actual wagers. In all four conditions of the task, the predicted wager significantly correlated with the number of points participants actually wagered: (i) advice stable phase r_1_ = 0.62, *p*_1_ = 3e-05; (ii) advice volatile phase r_2_ = 0.63, *p*_2_ = 2e-05; (iii) card stable phase r_3_ = 0.81, *p*_4_ = 9e-10; and (iv) card volatile phase r_4_ = 0.80, *p*_4_ = 1e-09; Figure S2). These findings suggest that the winning model captured variation in (the continuously measured) actual wager amount.

**Figure S2.**
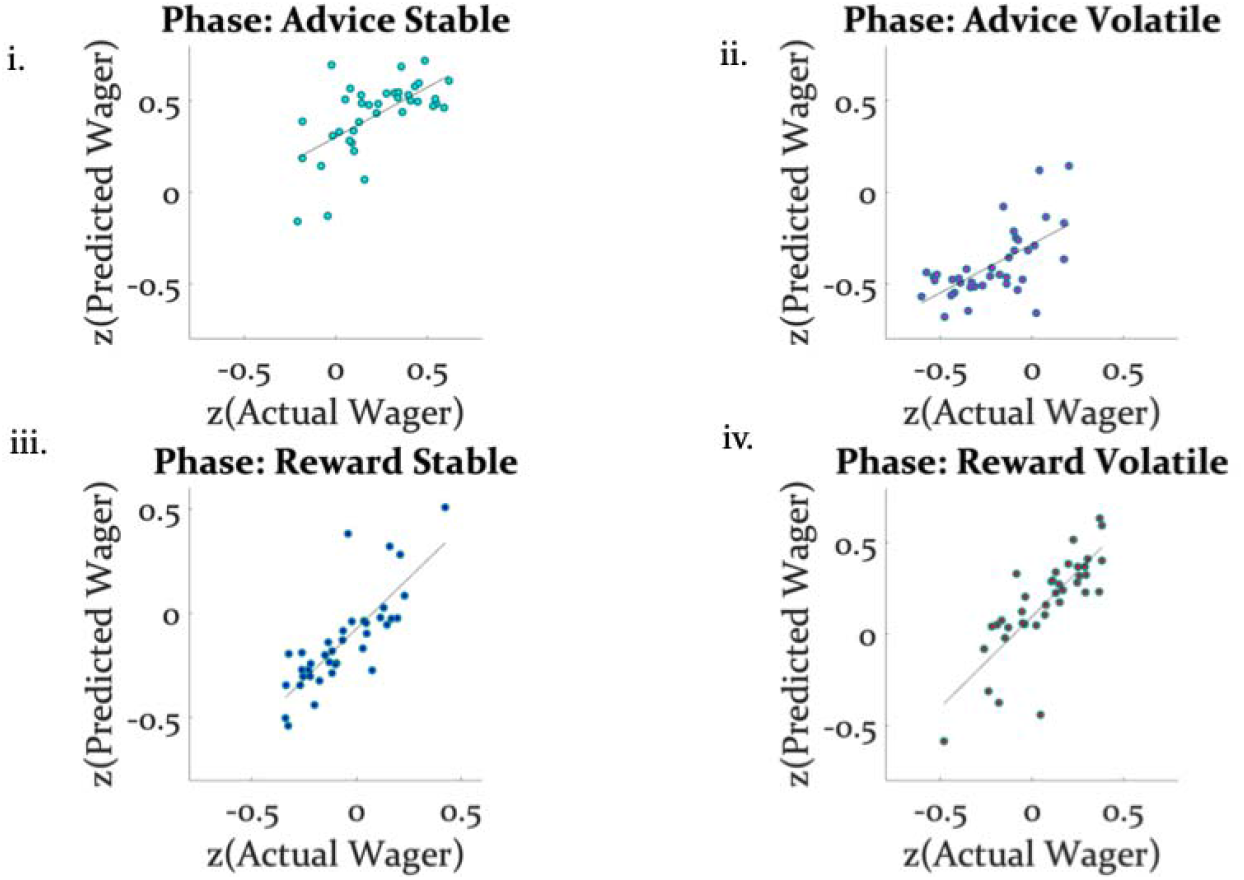
Model validity with regard to amount wagered: The z-transformed wager amount predicted by the model strongly correlated with the z-transformed number of points participants actually wagered across all four conditions of the task ((i) r_1_ = 0.62, *p*_1_ = 3e-05; (ii) r_2_ = 0.63, *p*_2_ = 2e-05; (iii) r_3_ = 0.81, *p*_4_ = 9e-10; (iv) r_4_ = 0.80, *p*_4_ = 1e-09). The regression line is plotted to illustrate the relationship between the actual and predicted wagers.

### Neural signatures of arbitration

Using behaviourally fitted computational trajectories to generate subject-specific GLMs for model-based fMRI analysis, we examined how the brain arbitrates between social and individual learning systems. We conceptualised the learning and arbitration process as hierarchical Bayesian inference, and fitted subject-specific trajectories reflecting arbitration (Eq. 20) to fMRI data.

Hierarchical precision-weighted PE signals were replicated in the same dopaminergic and frontoparietal regions as in previous studies using other sensory and social learning domains (see Iglesias et al., 2013; Diaconescu et al., 2017), indicating that the modifications in the experimental paradigm did not affect basic learning processes (see Supplementary material for the results).

Undirected tests for arbitration activity identified ventral prefrontal regions, such as the right ventromedial PFC (peak at [0, 46, −8]) and the right orbitofrontal cortex (OFC) [32, 30, −14]. Interestingly, frontal activations also included the right frontopolar cortex [4, 54, 30] and ventrolateral prefrontal cortex (VLPFC) [50, 36, 0], regions previously associated with arbitration between model-based and model-free forms of individual learning (Lee et al., 2014). The right VLPFC showing arbitration-related effects at [48, 35, −2] significantly overlapped with the arbitration-related reliability activations detected by Lee and colleagues, supporting the notion that arbitration is to some extent domain-independent.

More posterior regions showing effects of arbitration included occipital areas, the anterior insula, left thalamus, left putamen, bilateral middle cingulate sulcus, supplementary motor area (SMA) [−4, 0, 56], left dorsal middle cingulate gyrus [−10, −26, 44], the right amygdala [18, −10, −16] and the left midbrain [−6, −20, −10] (Table 3, Figure 6). Thus, a network of cortical and subcortical regions contributed to arbitration.

**Figure 6.**
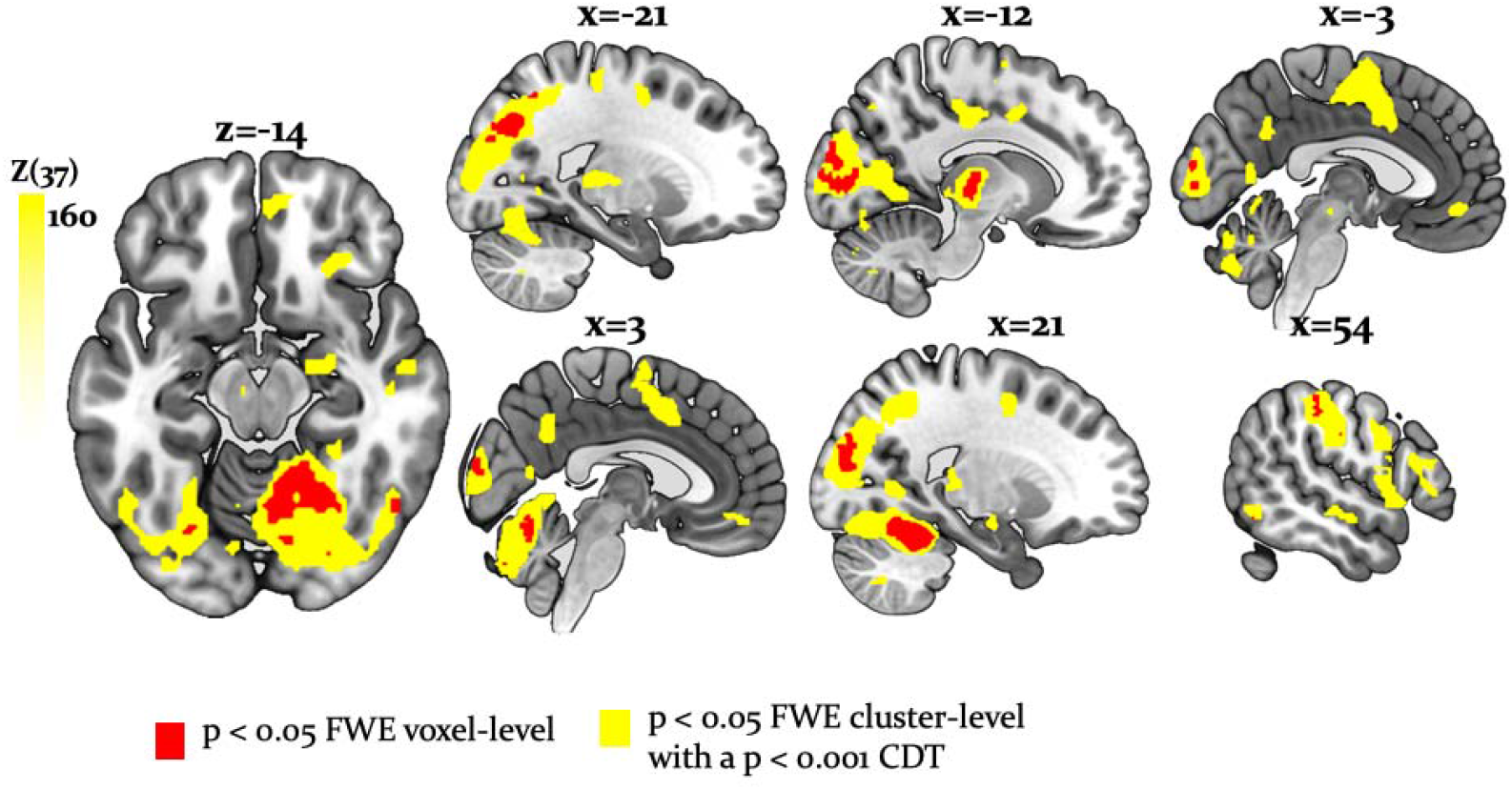
Whole-brain undirected arbitration signals: Effects of arbitration in favour of one or the other source of information were detected in ventromedial PFC, orbitofrontal cortex, right frontopolar cortex, VLPFC, the left midbrain, bilateral fusiform gyrus, lateral occipital gyrus, lingual gyrus, anterior insula, right amygdala, left thalamus, right cerebellum, bilateral middle cingulate sulcus and SMA. The figure shows whole-brain FWE voxel (red) – and cluster-level corrected (yellow) data, p < 0.05 (CDT = cluster defining voxel-level threshold). In this figure the scale reflects *F*-values.

**Table 3:**
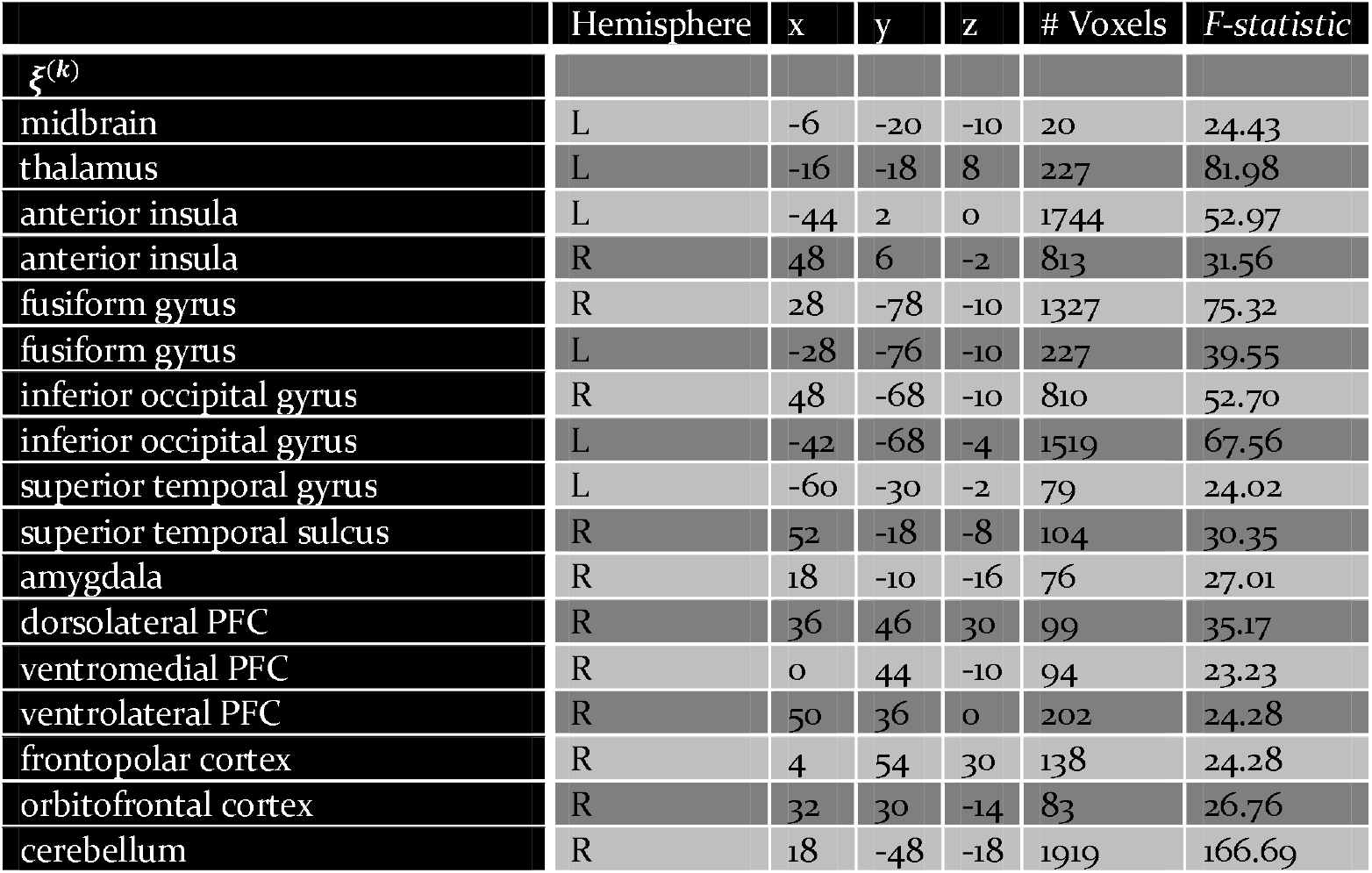
MNI coordinates and F-statistic of maxima of activations induced by either form of arbitration (Eqs. 19–20; p<0.05, cluster-level whole-brain FWE corrected). Related to Figure 6.

Directed tests for arbitration in favour of individual over social information identified activity increases in the right dorsolateral PFC [36, 46, 30], left SMA/anterior cingulate sulcus [−2, −8, 52] and the midbrain (Figure 7a). The BOLD signal change in these regions peaked during the time window of the wager decision. In summary, primarily dorsal regions of PFC were involved in arbitrating in favour of individually estimated card probability.

**Figure 7.**
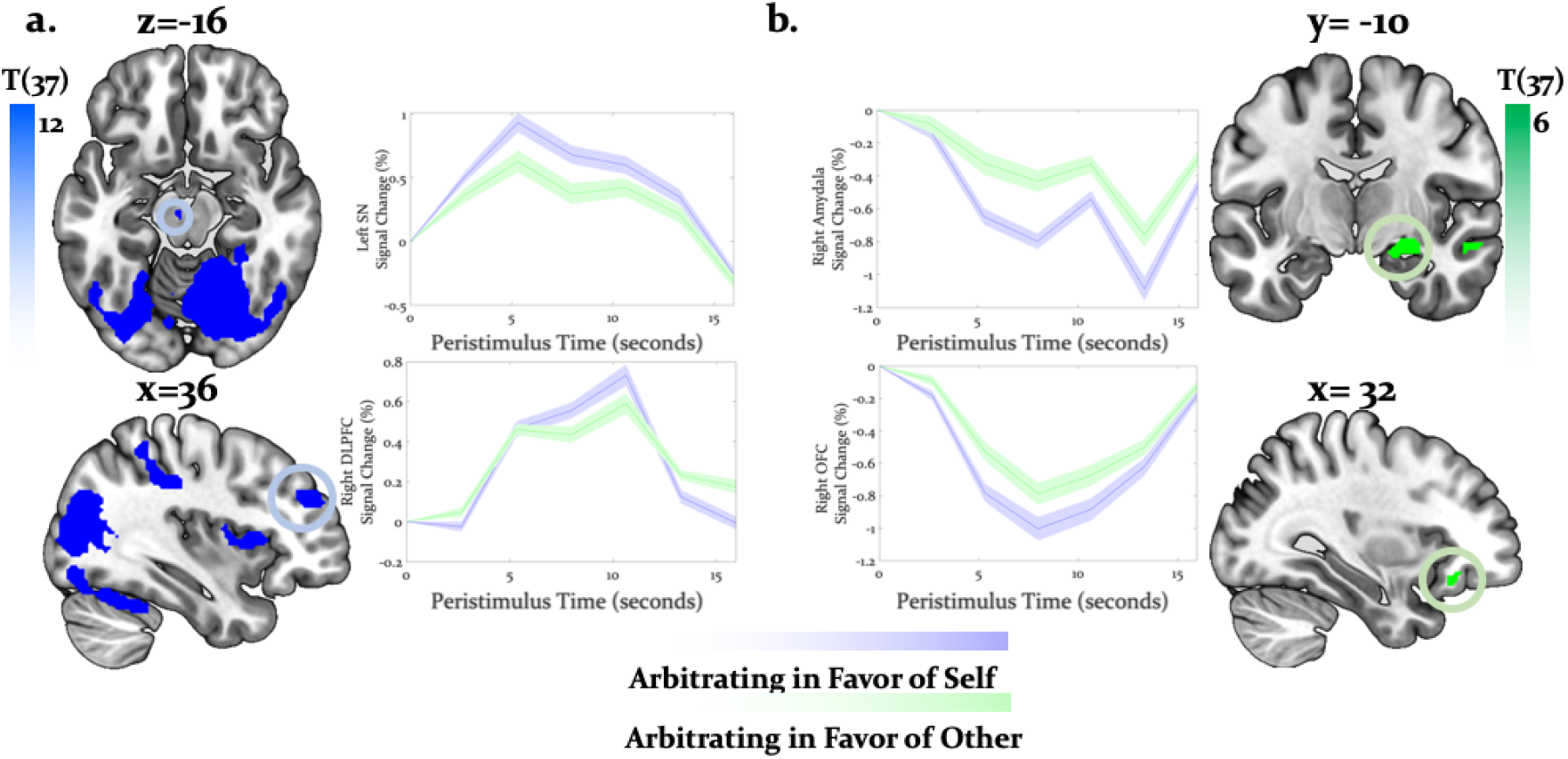
Neural arbitration directed to specific source of information: (a) Activity in the left midbrain (substantia nigra (SN)) [−6, −20, −10] (top) and the right DLPFC [36, 46, 30] (bottom) increased as participants arbitrated in favour of individually estimated card colour probability over the advisor’s suggestions when predicting the card colour (whole-brain FWE cluster-level corrected, p < 0.05). (b) Activity in right amygdala [18, −10, −16] (top) and right (OFC [32, 30, −14] (bottom) decreased to a lesser degree when participants arbitrated in favour of the advisor’s suggestion than when they arbitrated in favour of the individually learned estimates of card probability (whole-brain FWE cluster-level corrected, p < 0.05). The line plot reflects the average BOLD signal activity in the respective significantly activated cluster aligned to the onset of advice presentation relative to pre-advice baseline averaged across trials. Time series were averaged across trials for one representative subject in midbrain and DLPFC (a) or OFC and amygdala (b). The shaded areas depict +/- standard error of this mean. In this figure the scales reflect *t*-values.

Conversely, activity in the right amygdala, VLPFC, orbitofrontal and ventromedial PFC decreased as participants weighed the individually learned estimates of card probability over the advisor’s suggestion (Figure 7b). Outside PFC, the right anterior TPJ [56, −52, 24], right superior temporal gyrus [52, −18, −8], and right precuneus [6, −52, 32] showed similar effects (Tables 4 and 5 for the entire list of brain regions). Thus, primarily ventral regions of PFC together with temporal and parietal regions were more active during arbitration in favour of social information.

**Table 4:** MNI coordinates and t-statistic of maxima of activations induced by arbitration for the individually estimated card reward probability (Eq. 20; p<0.05, cluster-level whole-brain corrected). Related to Figure 7a.

|  | Hemisphere | x | y | z | # Voxels | t-statistic |
| --- | --- | --- | --- | --- | --- | --- |
| $\xi_c^{(k)}$: positive correlations | | | | | | |
| midbrain | L | -6 | -18 | -10 | 95 | 4.94 |
| thalamus | L | -16 | -18 | 8 | 232 | 5.10 |
|  | R | 22 | -30 | 4 | 206 | 5.10 |
| anterior insula | L | -44 | 2 | 0 | 2232 | 7.28 |
|  | R | 36 | 16 | 8 | 943 | 6.23 |
| supplementary motor area/<br>anterior cingulate sulcus | L | -2 | -8 | 52 | 1688 | 6.29 |
| dorsolateral PFC | R | 36 | 46 | 30 | 136 | 5.93 |
| middle occipital gyrus | R | 12 | -86 | 6 | 237 | 11.70 |
|  | L | -32 | -82 | 16 | 136 | 8.26 |
| superior occipital gyrus | R | 28 | -78 | 30 | 343 | 11.00 |
|  | L | -26 | -82 | 32 | 143 | 8.73 |
| cerebellum | R | 18 | -48 | -18 | 21557 | 12.91 |

**Table 5:** MNI coordinates and t-statistic of maxima of activations induced by arbitration for the social advice (Eq. 19; p<0.05, cluster-level whole-brain FWE corrected). Related to Figure 7b.

|  | Hemisphere | x | y | z | # Voxels | t -statistic |
| --- | --- | --- | --- | --- | --- | --- |
| $\xi_a^{(k)}$ : positive correlations | | | | | | |
| precuneus | R | 6 | -51 | 32 | 284 | 6.25 |
| amygdala | R | 18 | -10 | -16 | 107 | 5.20 |
| anterior cingulate cortex | L | -2 | 44 | -10 | 136 | 4.82 |
| ventromedial PFC | R | 8 | 52 | 14 | 231 | 5.72 |
| ventrolateral PFC | R | 50 | 36 | 0 | 305 | 4.93 |
| frontopolar cortex | R | 4 | 62 | 22 | 153 | 4.59 |
| orbitofrontal cortex | R | 28 | 26 | -16 | 126 | 5.11 |
| middle frontal gyrus | R | 38 | 14 | 28 | 305 | 5.36 |
| superior temporal gyrus | L | -60 | -30 | -2 | 107 | 4.90 |
| superior temporal sulcus | R | 52 | -18 | -8 | 152 | 5.51 |
| anterior temporoparietal junction | R | 56 | -52 | 24 | 173 | 4.18 |
| cerebellum | L | -24 | -84 | -34 | 121 | 4.11 |

To examine effects of arbitration in dopaminergic, cholinergic, and noradrenergic regions we also performed region-of-interest (ROI) analyses using a combined anatomical mask of dopaminergic, cholinergic, and noradrenergic nuclei. A single cluster in the right substantia nigra survived small-volume correction (p<0.05 FWE voxel-level corrected for the entire ROI; peak at [−6, −18, −10]; Figure 8). Activity in this region increased with arbitration in favour of individual estimates of card probabilities rather than advice.

**Figure 8.**
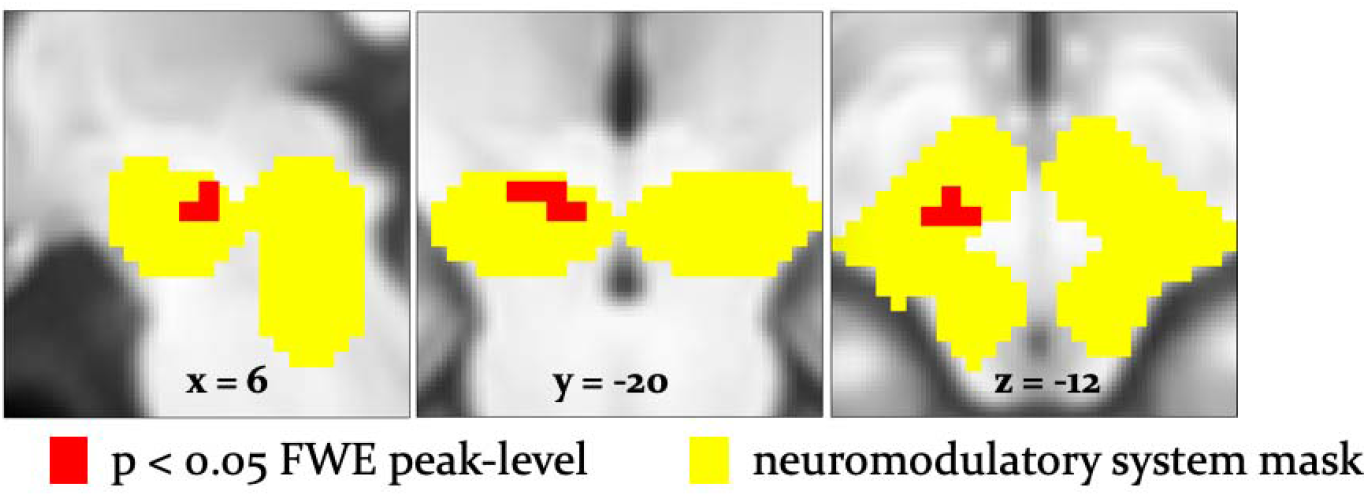
Arbitration signals in neuromodulatory ROI: Activation of the dopaminergic VTA/SN was associated with arbitrating in favour of individually learned information. Activation (red) is shown at p<0.05 FWE corrected for the full anatomical ROI comprising dopaminergic, cholinergic, and noradrenergic nuclei (yellow).

It is important to note that these regions showed significantly larger effects of arbitration than of the amount of points wagered. Responses reflecting arbitration dominated over responses reflecting wager magnitude in cerebellar, midbrain, occipital, parietal, ventral frontal, and temporal regions including the amygdala (Figure 9). In contrast, we observed stronger relation to wager magnitude in dorsolateral prefrontal cortex. As wager amount can be taken as a proxy for decision value or confidence (Lebreton et al., 2015), these data suggest that arbitration signals arise on top of decision value and confidence. Moreover, we captured arbitration as a model-derived, continuous, and time-resolved variable. Thus, our findings elucidate the process rather than the result of arbitration.

**Figure 9.**
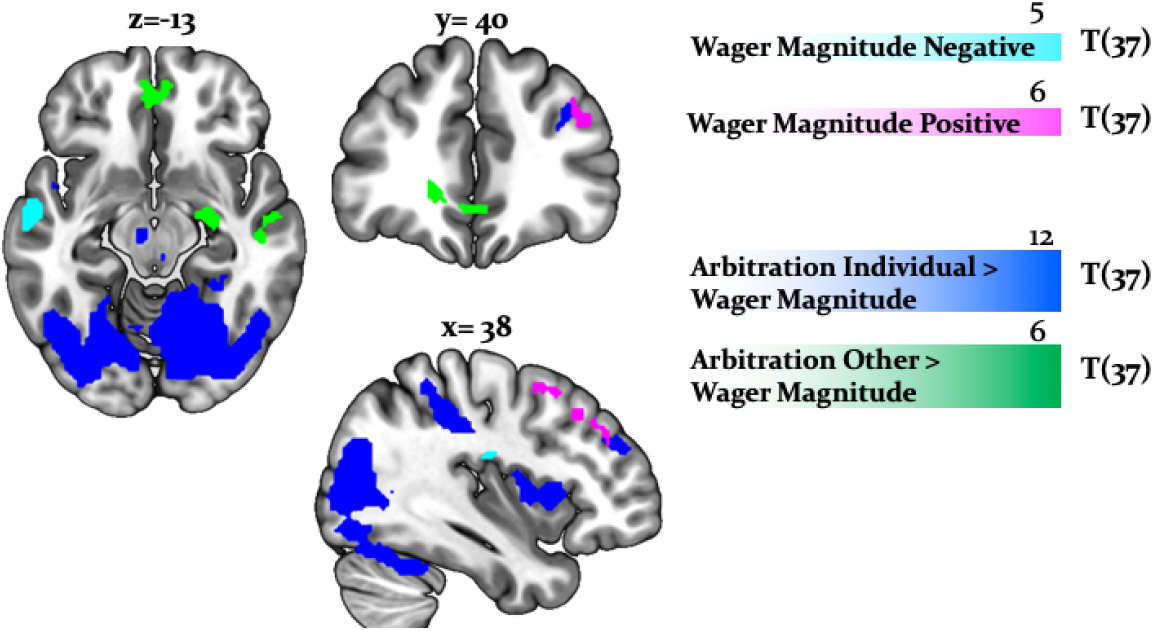
Arbitration vs. Wager Magnitude: Effects of arbitration (individual) (blue) were significantly larger in cortical and subcortical brain regions when compared to wager magnitude. Effects of arbitration in favour of social information were also significantly larger in ventromedial PFC and amygdala when compared to wager magnitude (green). Activity in middle frontal regions increased with increases in wager magnitude (magenta), whereas activity in the superior temporal gyrus decreased with increases in wager magnitude (cyan) (whole-brain FWE cluster-level corrected, p < 0.05).

### Main effect of stability and interaction with source of information

To examine arbitration from a different angle, we also conducted a factorial analysis. This was possible because we employed a 2 × 2 factorial design – i.e., two sources of information (individual versus social) in two different states (stable versus volatile) (Figure 10a). Specifically, we contrasted volatile with stable phases across both cue modalities. Volatility is closely tied to arbitration because it potentiates the perceived uncertainty associated with a given information source, and thereby the need to arbitrate. We assumed that arbitration increased as one of the two information sources was perceived as being more stable than the other. In all comparisons, we controlled for decision value and confidence by using the trial-wise wager amount as a parametric modulator in the analysis of brain data. We found two significant results (Figure 10): (i) a main effect of task phase (i.e., stability/volatility), and (ii) a significant interaction of task phase with source of information.

**Figure 10.**
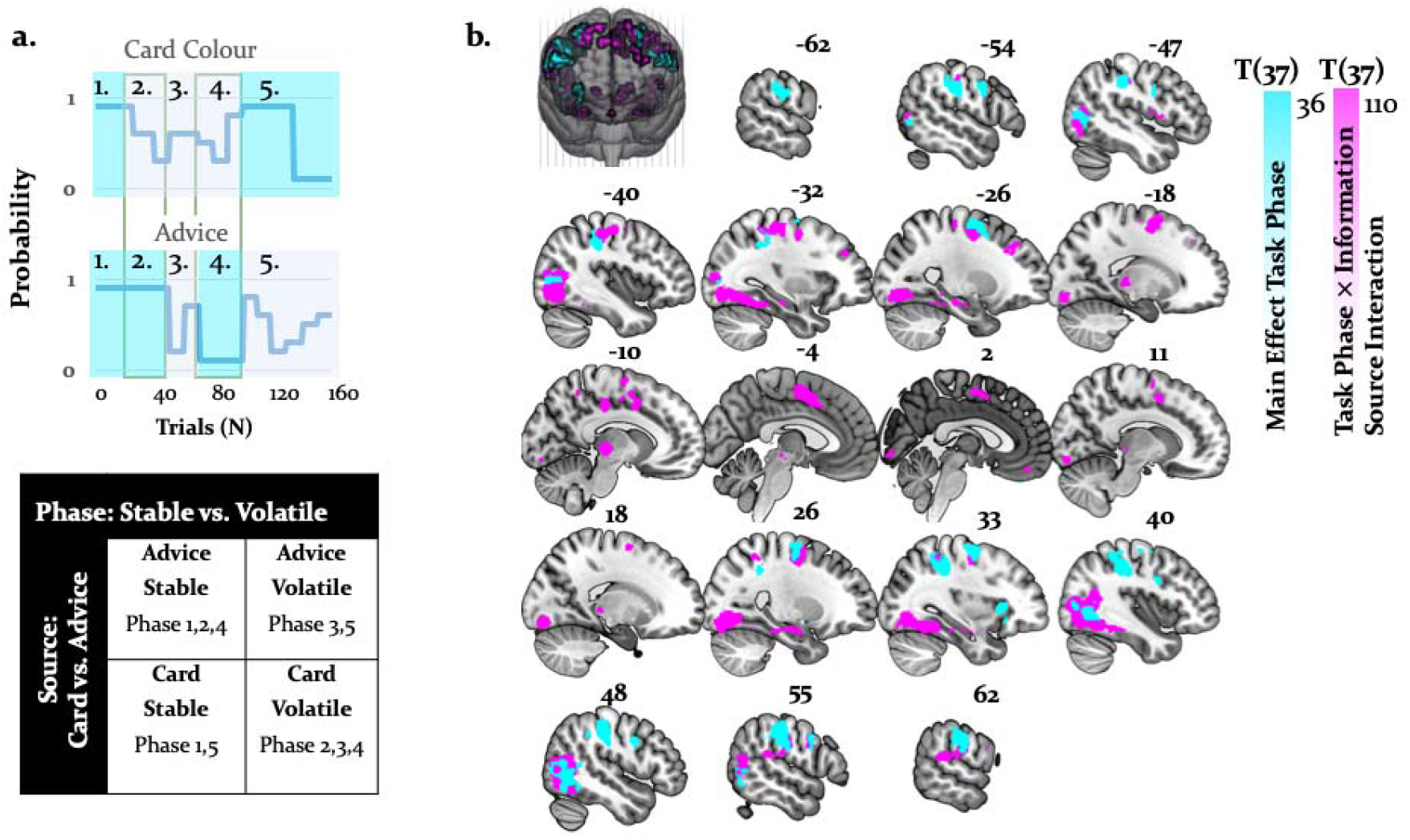
Activations related to task phase and interaction with source of information: (a) The task mapped onto a factorial structure with four conditions: (i) stable card and stable advisor, (ii) stable card and volatile advisor, (iii) volatile card and stable advisor, and (iv) volatile card and volatile advisor, as reflected by the shaded areas: yellow (stable), grey (volatile), (b) First, the main effect of stability irrespective of source of information activated primarily parietal regions and the anterior insula (cyan, whole-brain FWE cluster-level corrected, p<0.05). Second, the interaction between task phase and source of information was localized to left midbrain, occipital regions, anterior insula, thalamus, middle cingulate sulcus, SMA, OFC, and VLPFC (magenta, whole-brain FWE cluster-level corrected, p<0.05).

By contrasting stable against volatile phases, irrespective of information source, we found that the right supramarginal gyrus, bilateral inferior occipital gyrus, postcentral/precentral gyri, and the right anterior insula were more active for stable compared to volatile periods. Furthermore, an interaction between task phase and information source showed preferential activity for stable card information in the midbrain [−4, −22, −8]. Additional activations were detected in the right OFC, VLPFC, dorsal middle cingulate gyrus, and anterior cingulate sulcus/SMA (Figure 11, Tables 6 and 7). These regions processed stability (vs. volatility) more strongly for card than advice information. The regions processing stability (vs. volatility) more strongly for advice than card information also overlapped with the arbitration signal, and included the amygdala, the superior temporal sulcus, and the ventromedial PFC (Figure 11b). Thus, model-dependent and model-independent analyses agree in localizing arbitration to frontoparietal regions in the individual domain and to ventromedial prefrontal and amygdala regions in the social domain.

**Figure 11.**
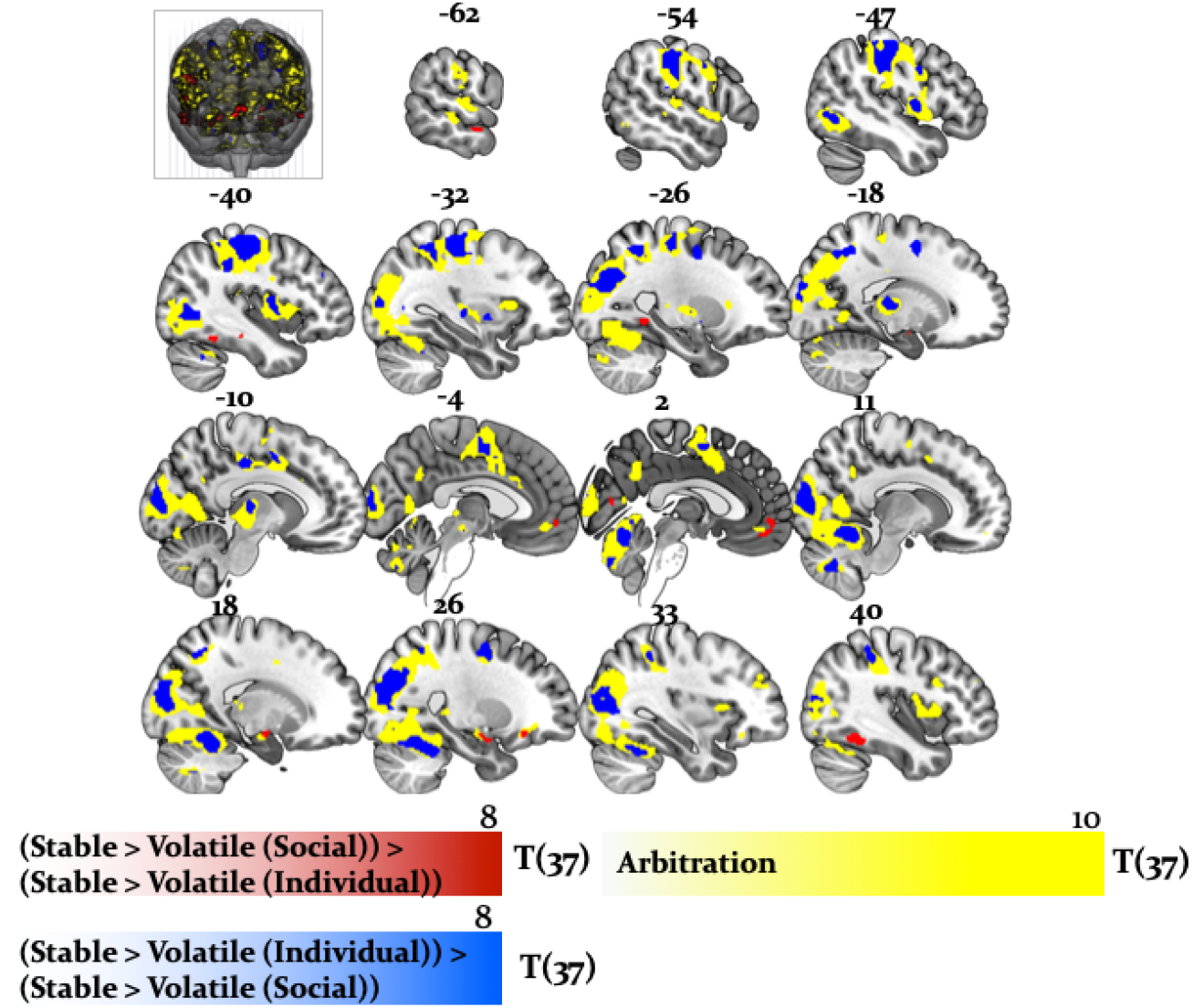
Overlap between model-dependent and model-independent results: Arbitration signal (Eqn. 19) (yellow) overlapped with the regions showing an enhanced effect of stability for individual compared to social learning systems (blue) and regions showing enhanced effects of stability in the social compared to individual learning systems (red) (whole-brain FWE peak-level corrected, p < 0.05).

**Table 6:** MNI coordinates and F-statistic for main effects of stability (p<0.05, FWE wholebrain corrected). Related to Figure 10 (activations in cyan).

|  | Hemisphere | x | y | z | # Voxels | F-statistic |
| --- | --- | --- | --- | --- | --- | --- |
| **Stability > Volatility** |  |  |  |  |  |  |
| supramarginal gyrus | R | 46 | -28 | 42 | 1199 | 38.16 |
| inferior occipital gyrus | R | 46 | -66 | 0 | 580 | 33.99 |
|  | L | -46 | -70 | 4 | 256 | 20.82 |
| anterior insula | R | 34 | 20 | 2 | 98 | 29.30 |
| postcentral gyrus | L | -52 | 2 | 34 | 107 | 28.97 |
|  | R | 54 | -22 | 34 | 129 | 5.59 |
| precentral gyrus | L | -60 | -20 | 32 | 512 | 40.21 |
|  | R | 50 | 4 | 32 | 129 | 20.58 |
| middle frontal gyrus | L | -26 | 0 | 58 | 117 | 20.18 |

**Table 7:**
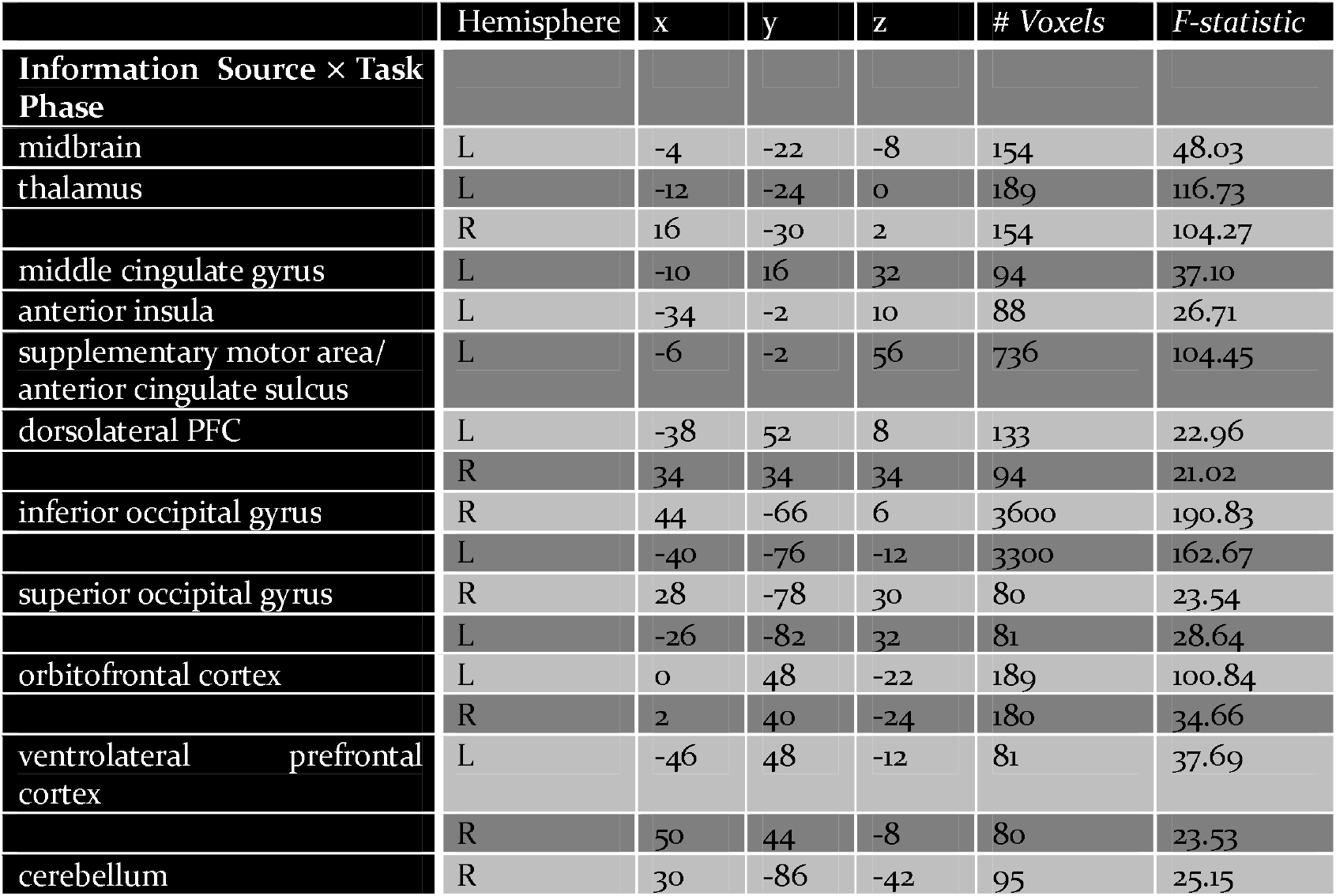
MNI coordinates and F-statistic for interactions between task phases and stimulus type (p<0.05, FWE whole-brain corrected). Related to Figure 10 (activations in magenta).

|  | Hemisphere | x | y | z | # Voxels | F-statistic |
| --- | --- | --- | --- | --- | --- | --- |
| **Information Source × Task Phase** |  |  |  |  |  |  |
| midbrain | L | -4 | -22 | -8 | 154 | 48.03 |
| thalamus | L | -12 | -24 | 0 | 189 | 116.73 |
|  | R | 16 | -30 | 2 | 154 | 104.27 |
| middle cingulate gyrus | L | -10 | 16 | 32 | 94 | 37.10 |
| anterior insula | L | -34 | -2 | 10 | 88 | 26.71 |
| supplementary motor area/<br>anterior cingulate sulcus | L | -6 | -2 | 56 | 736 | 104.45 |
| dorsolateral PFC | L | -38 | 52 | 8 | 133 | 22.96 |
|  | R | 34 | 34 | 34 | 94 | 21.02 |
| inferior occipital gyrus | R | 44 | -66 | 6 | 3600 | 190.83 |
|  | L | -40 | -76 | -12 | 3300 | 162.67 |
| superior occipital gyrus | R | 28 | -78 | 30 | 80 | 23.54 |
|  | L | -26 | -82 | 32 | 81 | 28.64 |
| orbitofrontal cortex | L | 0 | 48 | -22 | 189 | 100.84 |
|  | R | 2 | 40 | -24 | 180 | 34.66 |
| ventrolateral prefrontal cortex | L | -46 | 48 | -12 | 81 | 37.69 |
|  | R | 50 | 44 | -8 | 80 | 23.53 |
| cerebellum | R | 30 | -86 | -42 | 95 | 25.15 |

## Discussion

Our study shows how healthy subjects arbitrate between uncertain social and individual information under varying conditions of stability during a binary lottery task. (Figure 1). Participants arbitrated between the two information sources by taking into account their relative precision. The more precise one information source was over the other and the more stable the advisor was perceived to be, the more points participants were willing to wager.

By showing that participants tracked the volatility of both the advice and the card colour probabilities (Figure 3), our study underscores the importance of volatility in arbitrating between social advice and individual reward-relevant information. At the behavioural level, trial-by-trial accuracy of participant predictions, frequency of taking advice into account, and amount of points wagered on each trial (Figure S2) were all reduced by volatility. Thus, in stable compared to volatile environments, the propensity for arbitration in favour of the more precise information source increases. Numerous studies have demonstrated an important role of volatility in higher-level learning (Behrens et al., 2007, 2008; Nassar et al., 2010; Iglesias et al., 2013; Vossel et al., 2013; Diaconescu et al., 2017; Pulcu and Browning, 2017), in-keeping with the present findings.

Using both model-based and model-independent (factorial) fMRI analysis, we found that the arbitration signal correlated with activity in dorsolateral and ventrolateral PFC, frontopolar, and orbitofrontal cortex (Figure 6). These findings corroborate previous insights on arbitration between different forms of individual information also pointing to lateral prefrontal cortex (Lee et al., 2014), in line with domain generality for arbitrating. Note though that arbitration activity in prefrontal cortex followed a self-versus-other axis: dorsal prefrontal activity increased the more strongly participants weighed their own predictions of reward probabilities over the perceived reliability of the advisor. Conversely, activity in the ventromedial PFC and orbitofrontal cortex showed the opposite pattern and decreased in activity as participants relied more heavily on their own reward probability estimates relative to the advice (Figure 7). Together, arbitration appears to be sensitive to the source of information entering the arbitration process, contrary to an entirely domain-general process.

The results of both model-based and factorial analyses suggest a key role of the midbrain in arbitrating for individual estimates about card colour over advice (Figure 8). Primate studies found that sustained dopamine neuron activity signalled expected uncertainty (Fiorillo, 2003; Schultz, 2010; Schultz et al., 2008). This was further supported by human pharmacological studies (Burke et al., 2018; Ojala et al., 2018) as well as fMRI research showing possible involvement of dopamine in risk taking and of dopaminoceptive regions, such as the caudate, anterior insula, ACC and the medial PFC in uncertainty coding (e.g. (Dreher et al., 2006; Preuschoff et al., 2008; Tobler et al., 2009). In particular, studies employing hierarchical Bayesian models have identified VTA/SN activation correlated to precision of predictions about desired outcomes (Friston et al., 2014; Schwartenbeck et al., 2015).

These findings may also underscore the role of dopamine in modulating participants’ ability to optimise learning to suit ongoing estimates of environmental volatility. Potential neurobiological mechanisms include meta-learning models, which propose an important role of phasic dopamine signals in training the dynamics of the prefrontal system, to infer on the structure of the environment (Collins and Frank, 2016; Wang et al., 2018). Such models imply that improved learning of the structure of the environment, e.g. current levels of volatility, results in more appropriate arbitration adjustment.

The amygdala processed perceived reliability of social information, reflected in activity decreasing the more participants discounted the advisor relative to their own estimates of rewarded card colour probabilities. The amygdala has been implicated in processing facial expressions related to affective ToM (Schmitgen et al., 2016) and more generally, processing affective value and motivational significance of various stimuli, including other people (Güroğlu et al., 2008; Zink et al., 2008; Zerubavel et al., 2015). Together, the amygdala may represent the uncertainty of socially-relevant stimuli, inferred from intentions of others.

Similar to the amygdala, the orbitofrontal cortex showed a significant interaction between task phase and information source, indicative of arbitrating in favour of social information. This finding is consistent with the hypothesis that the orbitofrontal cortex and other areas of the social brain evolved to enable primates and particularly humans to successfully navigate complex social situations (Dunbar, 2009). This notion received support from strong positive correlations between orbitofrontal cortex grey matter volume and social network size (Powell et al., 2012), as well as sociocognitive abilities (Powell et al., 2010; Scheuerecker et al., 2010). Furthermore, in-keeping with a role of orbitofrontal cortex in mental state attribution for ambiguous social stimuli (Deuse et al., 2016), our findings suggest that this region reduces uncertainty of social cues signalling changes of intentionality.

An intriguing extension of the current study concerns the question whether arbitration occurs differently in patients with psychiatric and neurodevelopmental disorders involving ToM processes. If so, how and how do these processing differences affect behaviour? For example, individuals with autism spectrum disorder may preferentially rely on their own experiences rather than inferring predictions from the recommendations of others. Indeed, they appear to represent social prediction errors less strongly than individuals without autism (Balsters et al., 2017). Accordingly, they may be able to better infer the volatility of the card colour probability compared to the advice in our task. In contrast, patients with schizophrenia may be overly confident about their ability to judge advice validity due to fixed beliefs about the advisor’s intentions (Freeman and Garety, 2014) or show an over-reliance on social information in line with accounts of over-mentalization in this disorder (Montag et al., 2011; Andreou et al., 2015). Future work may test these intriguing possibilities.

## Limitations

One limitation of our study is that it did not include reciprocal social interactions, but rather used pre-recorded social partners. ToM processes may be more prominent in interactive paradigms (Diaconescu et al., 2014) or interactions that involve higher levels of recursive thinking (Devaine et al., 2014a, 2014b). By extension, our study may have limited generalizability to real-world social interactions. However, assessing arbitration between social and individual information necessitated the standardization of the advice given to each subject. To make the task as close as possible to a realistic social exchange, the videos of the advisor were extracted from trials when they truly intended to help or truly intended to mislead. More importantly, to adequately compare learning from social and individual information in stable and volatile phases, we needed to ensure that the two information types were orthogonal to each other and balanced in terms of volatility.

Second, we did not include a non-social control task. Accordingly, it is unclear how “social” the presently investigated form of learning about advisor fidelity and the volatility of advisor intentions is. To distinguish general inference processes under volatility from inference specific to intentionality, we previously included a control task (Diaconescu et al., 2014), in which the advisor was blindfolded and provided advice with cards from predefined decks that were probabilistically congruent to the actual card colour. This control task closely resembled the main task, with the exception of the role of intentionality. Model selection results suggested that participants did not incorporate time-varying estimates of volatility about the advisor into their decisions. In the current study, we tested this by including models without volatility, but found that they performed substantially worse than hierarchical models (see Figure 2 and Table 2a for details). Thus, our participants appeared to process advisor intentionality.

## Conclusions

Our study indicates that arbitrating between social and individual sources of information corresponds to weighing the relative reliability of each source. This process appears to engage different brain regions for social and individual information, in-keeping with domain specificity. However, the lateral prefrontal cortex appears to adjudicate between several different types of learning, in-keeping with domain generality. These findings contribute to our understanding of arbitration in neurotypical individuals, which may provide a knowledge basis for future insight into disorders with impaired arbitration.

## Materials and Methods

### Ethics Statement

The study was approved by the Ethics Committee of the Canton of Zürich. All participants gave written informed consent before taking part.

### Participants

We recruited 48 volunteers (mean age 23.6 ± 1.4, 32 females) who were non-smokers, right-handed, and had normal or corrected-to-normal vision. Participants had no history of neurological or psychiatric illness, or of drug abuse. Psychology students were excluded from participation because of previous exposure to similar advice-taking paradigms in their courses. Participants were asked to abstain from alcohol 24 hours prior to the study and from medication, including aspirin, 3 days prior to the study. We did not analyse the data of ten participants: two pilot participants; one participant who stopped the experiment midway due to head pain; one participant who fell asleep; and six participants where stimulus presentation malfunctioned during the experiment. Altogether, 38 participants (mean age 24.2 ± 1.3; 26 females) entered the final analysis.

### Stimuli and task

We modified the deception-free binary lottery game of Diaconescu and colleagues (2014). In each trial, the participant had to predict the colour of a card draw – blue or green. Participants could base their predictions on social information and/or on individually experienced recent outcome history (see below). They received social information from the “advisor”, who held up a card in one of the two colours before every draw, recommending to the participant which option to choose. The advisor based his or her suggestion on information that was true with a probability of 80%, although the participants were not informed of this fact. Furthermore, the advisor received monetary incentives to change his or her strategy and thus provide either helpful or misleading advice at different stages of the game (Figure 1b) with the average probability of advice being correct equivalent to 56%.

To display social information in a standardized fashion and gender-match advisors and participants, we created videos from two male and two female advisors, who changed their advice as a function of the incentives in a previously recorded face-to-face session (see Diaconescu et al, 2014). Their advice on each trial was recorded for an entire experimental session and the full-length videos were edited into 2-sec segments, focusing on the advice period. We received informed consent from all advisors in the initial (face-to-face) behavioural study to record and use the advice-giving videos in subsequent studies. All video clips were matched in terms of their luminance, contrast, and colour balance using Adobe Photoshop Premiere CS6. Advisor-to-participant assignment was randomized (within the gender-matching constraint) and balanced. We found no differences in performance and degree of reliance on advice between the four advisors: *F*(1,36) = 1.82, *p* = 0.16).

In contrast to previous studies (Diaconescu et al., 2014, 2017), participants had to infer card colour probabilities (blue versus green) from individually experienced outcomes of previous trials rather than being provided with (changing) pie charts explicitly stating the probabilities. In each trial they had to arbitrate between following either social information (previous advice, inferring on intention) or individual information (previous cards, inferring on probability). Moreover, also in contrast to previous studies, for each prediction participants wagered between one and ten points to indicate how confident they were about their predictions. The tick mark on the wager bar was randomly positioned in each trial to avoid providing a reference point (a regression analysis confirmed that the starting position of the wager indeed failed to explain each participant’s trialwise wager selection, t(37) = −0.89, p = 0.31). Depending on the correctness of the prediction, the wager was added to or subtracted from the cumulative score and thereby affected the participant’s payment at the end of the experiment (see below).

Each trial (Figure 1A) began with a video of the advisor holding up a card, followed by a decision screen in which participants selected the blue or green card. At the next screen, they were asked to provide the wager. The subsequent outcome screen revealed the drawn card. Finally, the updated cumulative score appeared. The colour-to-button assignment used to convey a decision (blue or green) and the orientation of the wager bar were randomized between participants to prevent confounding with visuomotor processes.

Across trials, the colour-reward probabilities and the advisor intentions varied independently of each other. In other words, the probability distributions of the two information sources – card colour and advice – were designed to be statistically independent. This allowed for a 2 × 2 factorial design structure, where trials could be divided into four conditions: (i) stable card and stable advisor, (ii) stable card and volatile advisor, (iii) volatile card and stable advisor, and (iv) volatile card and volatile advisor in a total of 160 trials (Figure 1B). Based on this factorial structure, we predicted that arbitration signals would vary as a function of the stability of each information source.

### Procedure

The actual experiment was divided into two sessions, with a two minute break in the middle when participants could close their eyes and rest. The first session included 70 trials and the second session 90 trials. We explained the deception-free task to participants and ensured their comprehension with a written questionnaire, which required them to describe the instructions in their own words.

To test the construct validity of our computational model and verify whether participants inferred on the advisor’s fidelity, we also asked them to rate the usefulness of the advisor’s card recommendation based on a multiple choice question (including, “helpful,”“misleading,” or “neutral”). This question was presented six times throughout the task and responses allowed us to assess whether at any point in time, the model could significantly predict participants’ responses.

Participants could earn a bonus of 10 Swiss Francs for a cumulative score of at least 380 points, and a bonus of 20 Swiss Francs for winning more than 600 points. Importantly, participants were not given any information about the bonus thresholds in order to prevent induction of local risk-seeking or risk-averse wagering behaviour (reference point effects) when participants were close to a threshold. Participants on average reached the first reward bonus and were paid 82.3 ± 8.4 Swiss Francs (including the performance-dependent bonus) at the end of the study. After the task, participants completed a debriefing questionnaire, and we revealed to them the general trajectory of the advisor’s intentions.

### Data Acquisition and Preprocessing

We acquired functional magnetic resonance images (fMRI) from a Philips Achieva 3T whole-body scanner with an 8-channel SENSE head coil (Philips Medical Systems, Best, The Netherlands) at the Laboratory for Social Neural Systems Research at the University Hospital Zurich. The task was presented on a display at the back of the scanner, which participants viewed using a mirror placed on top of the head coil. The first five volumes of each session were discarded to allow for magnetic saturation.

During the task, we acquired gradient echo T2*-weighted echo-planar imaging (EPI) data with blood-oxygen-level dependent (BOLD) contrast (slices/volume = 33; TR = 2665 ms; voxel volume = 2 × 2 × 3 mm^3^; interslice gap = 0.6 mm; field of view (FOV) = 192 × 192 × 180 mm; echo time (TE) = 35 ms; flip angle = 90°). The images were oblique, slices with −20° right-left angulation from a transverse orientation. The entire experiment comprised 1300 volumes, with 600 volumes in the first session and 700 in the second. Heart rate and breathing of the participants were recorded for physiological noise correction purposes using pneumatic belt and ECG.

We also measured the homogeneity of the magnetic field with a T1-weigh ted 3dimensional (3-D) fast gradient echo sequence (FOV = 192 × 192 × 135 mm^3^; voxel volume = 2 × 2 × 3 mm^3^; flip angle = 6°; TR = 8.3 ms; TE1 =2 ms; TE2 = 4.3 ms). After the experiment, we acquired T1-weighted structural scans from each participant using an inversion-recovery 3-D fast gradient echo sequence (FOV = 256 × 256 × 181 mm^3^; voxel volume = 1 × 1 × 1 mm^3^; TR = 8.3 ms; TE = 3.9 ms; flip angle= 8°).

The software package SPM12 version 6470 (Wellcome Trust Centre for Neuroimaging, London, UK; http://www.fil.ion.ucl.ac.uk/spm) was used to analyse the fMRI data. Temporal and spatial preprocessing included slice-timing correction, realignment to the mean image, and co-registration to the participant’s own structural scan. The structural image underwent a unified segmentation procedure combining segmentation, bias correction, and spatial normalization (Ashburner and Friston, 2005); the same normalization parameters were then applied to the EPI images. As a final step, EPI images were smoothed with a Gaussian kernel of 6mm full-width half-maximum.

BOLD signal fluctuations due to physiological noise were modelled with the PhysIO toolbox (http://www.translationalneuromodeling.org/tapas) (Kasper et al., 2017) using Fourier expansions of different order for the estimated phases of cardiac pulsation (3rd order), respiration (4th order) and cardio-respiratory interactions (1st order; (Glover et al., 2000)). The 18 modelled physiological regressors entering the subject-level GLM along with the six rigid-body realignment parameters and regressors of interest were used to account for BOLD signal fluctuations induced by cardiac pulsation, respiration, and the interaction between the two.

### Computational Modelling

We formalised arbitration in terms of hierarchical Bayesian inference as the relative perceived reliability of each information source. In other words, arbitration was defined as a ratio of precisions: the precision of the prediction about advice accuracy and colour probability, divided by the total precision. The precisions of the predictions afforded by each learning system are obtained by applying a two-branch hierarchical Gaussian filter (Mathys et al., 2011, 2014) along with a response model (see below) to participants’ trial--wise behaviour (i.e., choices and wagers).

#### Learning Model: Hierarchical Gaussian Filter

The HGF is a model of hierarchical Bayesian inference widely used for computational analyses of behaviour (e.g., (Iglesias et al., 2013; Vossel et al., 2013; Hauser et al., 2014; de Berker et al., 2016; Marshall et al., 2016). To apply it to our task, we assumed that the rewarded card colour (individual learning) and the advice accuracy (social learning) varied as a function of hierarchically coupled hidden states: 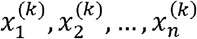. They evolved in time by performing Gaussian random walks. At every level, the step size was controlled by the state of the next-higher level (Figure 3A).

Starting from the bottom of the hierarchy, states *x*_1,*a*_ and *x*_1,*c*_ represented binary (0 or 1) variables, namely the advice accuracy (1 for accurate, 0 for inaccurate) and the rewarded card colour (1 for blue, 0 for green). All states higher than *x*_1_ were continuous. They denoted (i) the advisor fidelity and tendency for a given card colour to be rewarded, and (ii) the rate of change of the advisor’s intentions and card colour contingencies, respectively. Four learning parameters, namely, *κ_a_*, *κ_c_*, *ϑ_a_* and *ϑ_c_* determined how quickly the hidden states evolved in time. Parameter *κ* represented the degree of coupling between the second and the third levels in the hierarchy, whereas *ϑ* determined the variability of the volatility over time (meta-volatility). This constitutes the *generative model* of the process producing the outcomes observed by subjects. The overall model and the formal equations describing these relations in a social learning context are detailed in Diaconescu et al., 2014.

#### Model Inversion: Agent-specific arbitration

In accordance with Bayes’ rule, we assumed that participants who make inferences on advice and card colours form posterior beliefs over the hidden states (i.e., congruency of advice with actual card colour; rewarded card colour) based on the outcomes they observe. Model inversion is the application of Bayes’ rule to a generative model such as the one described above. This leads to a *recognition* or *perceptual model*, which describes subjects’ beliefs about hidden states. Assuming Gaussian distributions, these agent-specific beliefs are denoted by their summary statistics, i.e., *μ* (mean) and *σ* (variance/uncertainty) or the inverse of the variance *π* = 1/*σ* (precision/certainty).

Using variational Bayes under the mean-field approximation, simple analytical trial-by-trial update equations can be derived. The posterior means 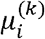 or predictions on each trial *k* at each level of the hierarchy *i* change as a function of precision-weighted prediction errors (PEs):

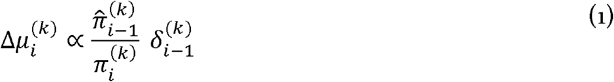

Throughout, predictions or prior beliefs about the hidden states (before observing the outcome) are denoted with a hat symbol. States 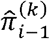 and 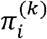 represent the estimated precisions about (i) the input from the level below (i.e., precision of the data – advice congruency or rewarded card colour) and (ii) the belief at the current level, respectively.

The updates about the advisor’s fidelity are:

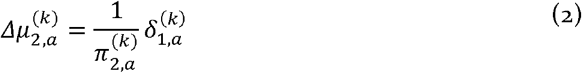

where

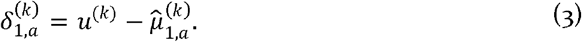

Variable *u*^(*k*)^ is the sensory input at trial *k*, where given advice is either accurate (*u*^(*k*)^ = 1) or inaccurate (*u*^(*k*)^ = 0). Furthermore, 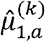 corresponds to the logistic sigmoid of the current expectation of the advisor fidelity:

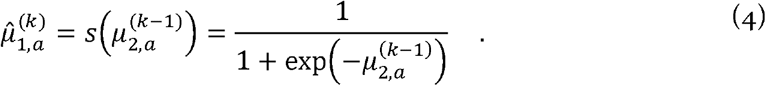

The current belief precision is equivalent to:

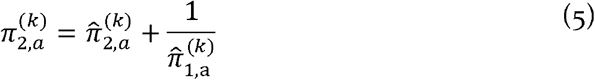

with the predicted (i) belief precision 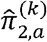 and (ii) the sensory, lower-level precision about the advice 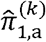 computed as:

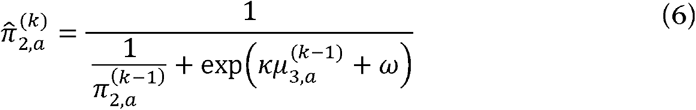

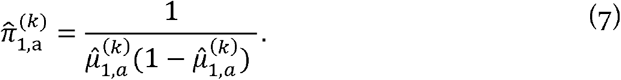

Thus, the advice belief precision depends on (i) the predicted sensory precision of the input, 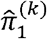 and (ii) the predicted volatility, 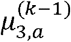 from the level above via equation 6.

The precision-weighted PE about the advice, which is used to update the belief about fidelity is equivalent to:

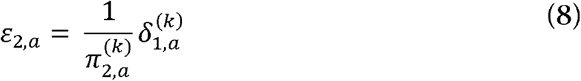

Going up the hierarchy, the updates of advice volatility are proportional to precision-weighted PEs:

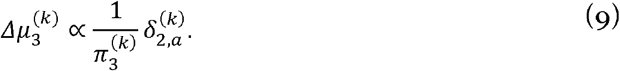

They depend on the higher-level volatility PE *δ*_2,*a*_:

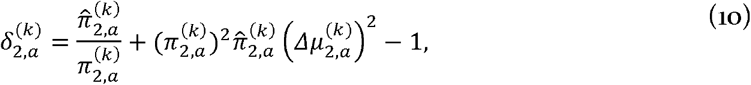

and the higher-level volatility precision *π*_3_:

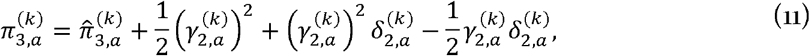

with the precision of the prediction about volatility given by

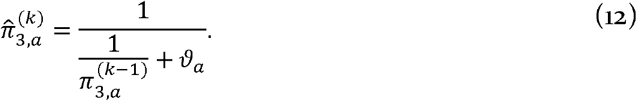

The third level, the precision-weighted volatility PE is equivalent to:

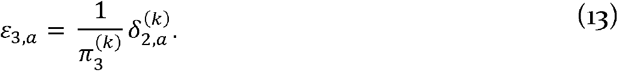

The same form of update equations (and precision-weighted PEs) can be derived for the individual information source, updating beliefs about the rewarded card colour, i.e.:

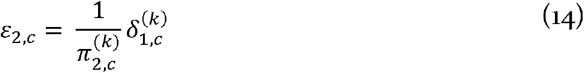

and

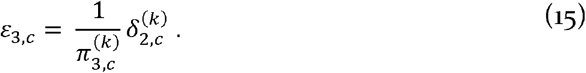

The prediction errors exhibit a similar form as for the advice, with

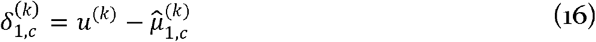

for the outcome PE and

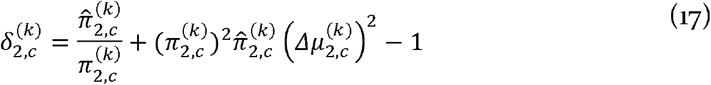

for the card volatility PE. The individually estimated card colour probability is equivalent to the logistic sigmoid of the current expectation of the rewarding card colour:

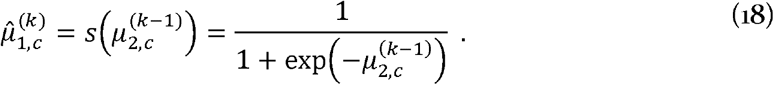

In this context, Bayes-optimality is individualized with respect to the values of the learning parameters, which were allowed to differ across subjects.

##### Arbitration Signal

Within this computational framework, we defined arbitration as the relative perceived precision associated with each information source, which is equivalent to the precision of the prediction of each information channel (advice or card; i.e., 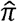) divided by the total precision. Arbitration is consistent with Bayes’ rule representing the optimal integration of the two inferred states by their precisions.

Arbitration towards advice – i.e., the perceived reliability of the social information source is equivalent to:

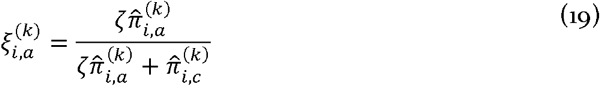

on each trial *k* at each level of the hierarchy *i* with *ζ* as the social bias or the additional bias towards the advice.

At the first level and at *i* = 1, the participant relies preferentially on the social input during action selection when 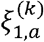 exceeds 0.5. Conversely, when 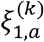 is below 0.5 (see Eq. 9), the participant relies more on individual (estimates of) card colour probabilities:

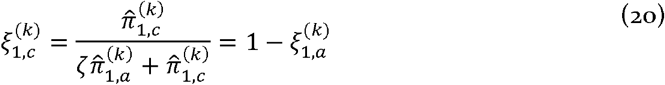

#### Response Model

To map beliefs to decisions, we assumed that the prediction of card colour on a given trial *k* is a function of arbitration and of the predictions afforded by each source (see Eq. 21). The response model predicts two components of the behavioural response: (i) the participant’s decision to accept or reject the advice and (ii) the number of points wagered on every trial. Responses were coded as *y* = 1 when participants took the advice and chose the card colour indicated by the advisor, and *y* = 0 when participants decided against following the advice and chose the opposite card colour. The expected outcome probability is thus a precision-weighted sum of the two information sources, the estimates of advice accuracy and rewarding colour probability.

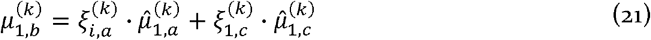

where 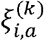 and 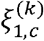 are the arbitration for each information source; 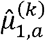 is the expected advice accuracy (Eq. 4) and 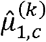 is the transformed expected card colour probability from the perspective of the advice (i.e., the estimated card colour probability indicated by the advisor).

The probability that participants chose a particular card colour according to their expectations about the outcome (Equation 21) was modelled by a softmax function:

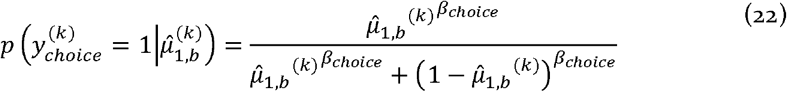

where *β_choice_* > 0 is the subject-specific inverse decision temperature parameter. A low decision temperature (high *β_choice_*) means always choosing the highest probability colour, whereas a high decision temperature (low *β_choice_*) means sampling randomly from a uniform distribution.

The trial-wise wager response was formalised as a linear function of various sources of uncertainty/precision: (i) irreducible decision uncertainty or 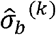 about the outcome, (ii) arbitration, (iii) informational uncertainty about the card colour or the advice, and (iv) environmental uncertainty/volatility about the card colour or the advice. We transformed these computational quantities down to the first level in the hierarchy using the sigmoid transformation and used them to predict the trial-by-trial wager (Figure 5 for the group average of each of these quantities):

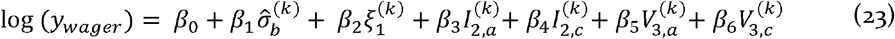

with

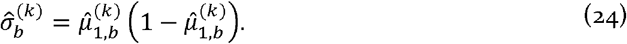

Parameter *ζ* captures the social bias in arbitration (equation 19) and 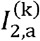 is the informational uncertainty about the advisor fidelity

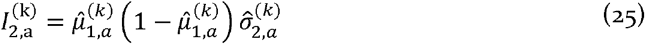

where 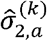 is the inverse of 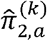 and represents the informational uncertainty of the prediction about the advisor’s fidelity (Eqn. 6).

The environmental volatility is defined as:

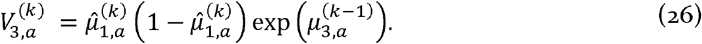

Equivalent equations can be derived for the individual information source.

The trial-wise wager magnitude predicted by the model is then defined as:

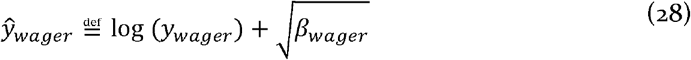

where *β_wager_* is a stochasticity parameter associated with the wager amount. For the priors of all *β* parameters estimated here, please refer to Table 1.

### Competing Models

We considered two families of perceptual models. The first family included the full, three-level version of the HGF (as described above). By contrast, the second family lacked the third level, and assumed that agents do not estimate the volatility of the card probabilities or the advice. Thus, comparing families with and without volatility tested whether volatility mattered for arbitrated behaviour.

For the response model, we considered three families, capturing different ways in which participants may arbitrate between social and individual sources of information to make decisions. These included: (i) an “Arbitrated” model, which assumed that participants combine and arbitrate between the two information sources, possibly unequally, (ii) an “Advice only” model, assuming arbitration-free reliance on social information only, and (iv) a “Card only” model, representing arbitration-free reliance on the inferred card colour probabilities only (Figure 2a).

All models were compared formally using Bayesian model selection (BMS; (Stephan et al., 2009). Random effects BMS results in a posterior probability for each model given the participants’ data. The relative goodness of models is denoted by the “protected exceedance probability” reflecting how likely it is that a given model has a higher posterior probability than any other model in the set of models considered (Stephan et al., 2009; Rigoux et al., 2014).

We adopted a similar set of priors over the perceptual model parameters as in our previous studies (Diaconescu et al., 2014) (see Table 1). Maximum-a-posteriori (MAP) estimates of model parameters were obtained using the HGF toolbox version 3.0, freely available as part of the open source software package TAPAS at http://www.translationalneuromodeling.org/tapas.

### FMRI Data Analysis

#### Single-subject Level

Our fMRI data analysis focused on the neural mechanisms of arbitration. Specifically, we conducted two types of analyses on the pre-processed fMRI data:

First, we performed a model-based fMRI analysis, in which we constructed a general linear model (GLM), which sought to explain the high-pass filtered voxel time-series with several parametric modulators. The parametric modulators are listed below and were derived from the winning model (i.e., arbitrated three-level version of the HGF, which had the highest posterior probability at the group level). The GLMs were individualized, as the regressors were obtained from fitting the model to the behavioural data of each of the 38 participants. We individualized GLMs because participants differed in how much they relied on each information source and in the extent to which volatility influenced their trial-by-trial wagers (Figures S1). To investigate the unique contribution of each parametric modulator, we did not orthogonalise them. Moreover, we also included movement and the physiological noise regressors obtained from the PhysIO toolbox (Kasper et al., 2017) based on ECG and respiration recordings as regressors of no interest.

In addition to arbitration at the time of advice presentation, we modelled the wager and the outcome phases to examine the effects of hierarchical precision-weighted PEs, and thus test the validity of the computational model and the reproducibility of previous findings, see Supplementary material (Iglesias et al., 2013; Diaconescu et al., 2017). Specifically, the following regressors were included in the GLM:

1. **Cue presentation** – time when the advice was presented (regressor duration 2s);
2. **Arbitration** – parametric modulator of (1), using the trial-specific arbitration quantity (Eq. 19–20);
3. **Wager presentation** – time when the option to wager was presented (regressor duration 0s);
4. **Wager** – parametric modulator of (3), using the trial-specific amount of points wagered;
5. **Outcome** – time when the winning card colour was presented (regressor duration 0s);
6. **Advice Precision-weighted PE** – parametric modulator of (5), using the trial-specific precision-weighted PE of advice validity (Eq. 8);
7. **Outcome Precision-weighted PE** – parametric modulator of (5), using the trial-specific precision-weighted PE arising from comparing actual and predicted card colour (Eq. 14).
8. **Volatility Advisor Precision-weighted PE** – parametric modulator of (5), using the trial-specific precision-weighted PE of advice volatility (Eq. 13);
9. **Volatility Card Precision-weighted PE** – parametric modulator of (5), using the trial-specific precision-weighted PE of card colour volatility (see Eq. 15).

We observed no significant correlations between response times (RTs) and any of the parametric modulators (|r|<0.3, p>0.05) and therefore did not model RT explicitly. The lack of effects on RTs may be due to the temporal structure of our task (Figure 1). Specifically, participants responded long after having received individual information (card outcome in previous trial) and social information had fixed duration (video). Therefore, they are likely to have simply conveyed the decision in the response phase but made it at some time during the video or even before.

Second, we predicted that arbitration should be sensitive to volatility, and favour one or the other source of information as a function of perceived relative reliability. Based on this hypothesis, we also performed a non-model based, factorial analysis by dividing the 160 trials into four conditions: (i) stable card and stable advisor, (ii) stable card and volatile advisor, (iii) volatile card and stable advisor, and (iv) volatile card and volatile advisor (Figure 10a). This GLM included for each of the four conditions the time when the advice was presented (the cue phase) and the trial-wise amount wagered as a parametric modulator. We assumed that the difference between the four conditions will be expressed in the advice (cue) phase, before subjects make their predictions.

#### Group Level

Contrast images from the 38 participants entered a random effects group analysis (Penny and Holmes, 2007). We used F-tests to identify undirected arbitration signals. Moreover, one-sample t-tests to investigate directed social or individual arbitration signals and positive or negative BOLD responses for each of the computational trajectories of interest described above.

Participant gender and age were included as covariates of no interest at the group level (the findings remained the same without these covariates). To investigate individual variability in the representation of social arbitration as a function of reliance on advice, we used parameter *ζ* to perform a median split of the group of participants.

For all analyses, we report results that survived whole-brain family-wise error (FWE) correction at the cluster level at p < 0.05, under a cluster-defining threshold of p < 0.001 at the voxel level using Gaussian random field theory (Worsley et al., 1996). Given recent debate regarding the vulnerabilities of cluster-level FWE procedures (Eklund et al., 2016), it is worth emphasising that this cluster-defining threshold ensures adequate control of cluster-level FWE rates in SPM (Flandin and Friston, 2016). The coordinates of all brain regions were expressed in Montreal Neurological Institute (MNI) space.

Based on recent results that precisions at different levels of a computational hierarchy may be encoded by distinct neuromodulatory systems (Payzan-LeNestour et al., 2013; Schwartenbeck et al., 2015), we also performed region-of-interest (ROI) analyses based on anatomical masks. We included (i) the dopaminergic midbrain nuclei substantia nigra (SN) and ventral tegmental area (VTA) using an anatomical atlas based on magnetization transfer weighted structural MR images (Bunzeck and Düzel, 2006), (ii) the cholinergic nuclei in the basal forebrain and the tegmentum of the brainstem using the anatomical toolbox in SPM12 with anatomical landmarks from the literature (Naidich and Duvernoy, 2009) and (iii) the noradrenergic locus coeruleus based on a probabilistic map (Keren et al., 2009).

The routines for all analyses performed here are publicly available as Matlab code: https://github.com/andreeadiaconescu/wagad.

## Acknowledgments

We are grateful for support by the Swiss National Science Foundation (Ambizione grant PZ00P3_167952 to AOD; PP00P1_150739, 100014_165884, and 100019_176016 to PNT) and the Krembil Foundation to AOD. We are also grateful to Klaas Enno Stephan for providing guidance and funding for the study.

