## Supplementary Material and Tables for "Neural Arbitration between Social and Individual Learning Systems"

Correspondence should be addressed to:

Andreea O. Diaconescu, PhD

Krembil Centre for Neuroinformatics (CAMH)

250 College St. M5T 1R8

#### Replication of Hierarchical Precision-weighted PE Effects across Learning Domains

To test whether the current task study replicates previous findings on the representation of hierarchical precision-weighted PEs (Iglesias et al., 2013; Diaconescu et al., 2017), we performed the same model-based analysis using Bayesian surprise (equivalent to an unsigned precision-weighted outcome PE; the absolute value of equation 14). Replicating the previous study (Iglesias et al., 2013), we found that the outcome-related BOLD activity of the substantia nigra positively correlated with the unsigned precision-weighted outcome PE, as did the anterior insula, (ventro)lateral PFC, and the intraparietal sulcus (Figure S3 and Table S1). In the previous study, participants predicted a visual outcome using an auditory cue (Iglesias et al., 2013). Thus, the PE coding of these regions seems to be modality-independent.

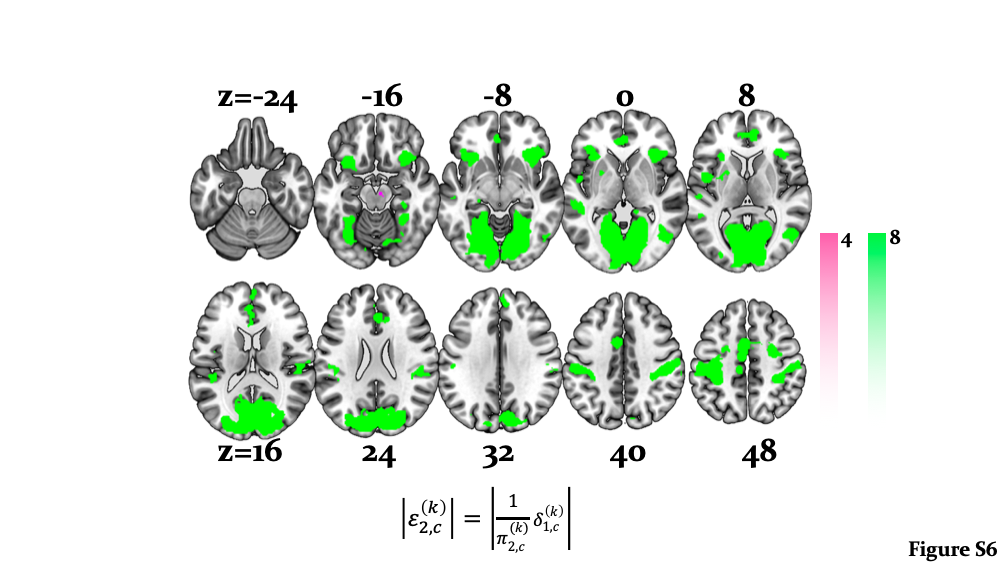

Figure S3| **Main effects of precision-weighted PEs in individually estimated card colour probabilities** (equation 14)**:** Whole-brain activation by $\varepsilon_{2,c}$ (**green**): Activations by unsigned precision-weighted PE about the card colour probabilities were detected in the bilateral inferior/middle occipital gyri, anterior insula, bilateral inferior, medial and middle frontal gyri, and the bilateral intraparietal sulcus (whole-brain FWE voxel- and cluster-level corrected, p<0.05). Activation of the right substantia nigra associated with the unsigned precision-weighted prediction error about the card colour probabilities is highlighted in **magenta**. This activation is shown at p<0.05 FWE corrected for the volume of our anatomical mask comprising dopaminergic nuclei.

With respect to the signed precision-weighted advice PE (equation 8), we also reproduced results from a recent study that employed a different advice-taking paradigm, where participants learned about advice and integrated it along with unambiguous individual information to predict the outcome of a binary lottery (Diaconescu et al., 2017). Effects of signed precision-weighted advice PE were detected in right VTA/substantia nigra, the right insula, left middle temporal cortex, right dorsolateral, left dorsomedial and middle frontal cortex (Figure S4 and Table S2).

Please note that we used the unsigned (absolute) precision-weighted PEs for the card outcomes, but the signed precision-weighted PEs for the advice. In the case of the card, the sign of this PE depends on an arbitrarily chosen coding of the colour and the sign is meaningless (see Iglesias et al., 2013). In contrast, for the advice the sign refers to the valence and instances of surprise where the advisor was more helpful than predicted may have a different meaning than instances of surprise where the advisor was more misleading than predicted (see Diaconescu et al., 2017). For completeness, we also investigated the neural correlates of the signed reward precision-weighted PE and noted a similar network of posterior parietal and dorsolateral prefrontal regions.

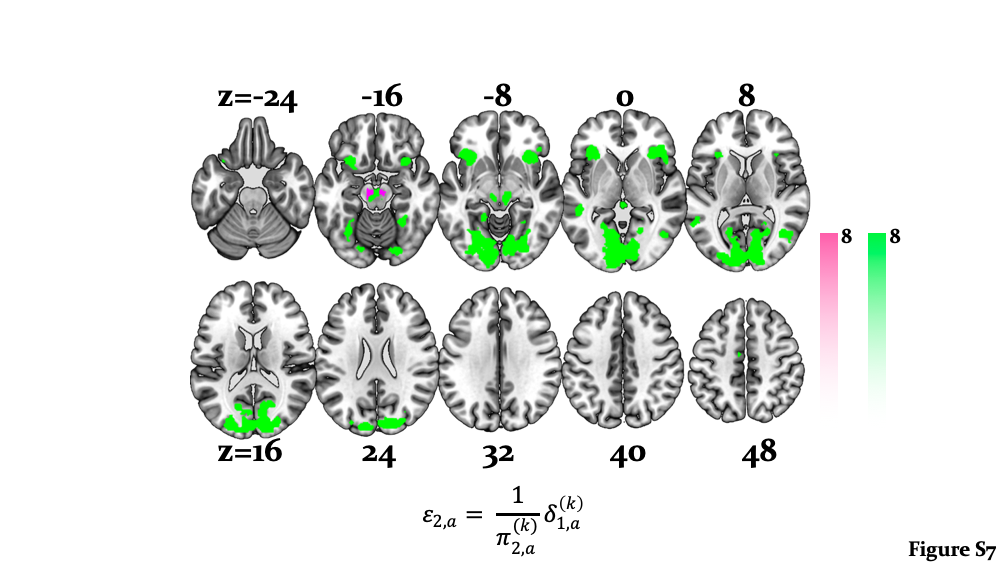

Figure S4| **Main effects of precision-weighted PEs about advisor fidelity** (equation 8): Whole-brain activation by $\varepsilon_{2}$ advice (**green**): Activations by signed precision-weighted PE about advisor fidelity were detected in the bilateral fusiform gyrus, lingual gyrus, anterior insula, left middle temporal cortex, hippocampus, right temporal-parietal junction, bilateral dorsolateral and left dorsomedial prefrontal cortex (whole-brain FWE voxel- and cluster-level corrected, p<0.05). Activation of the bilateral VTA associated with the signed precision-weighted prediction error about advisor fidelity is highlighted in **magenta**. This activation is shown at p<0.05 FWE corrected for the volume of our anatomical mask comprising dopaminergic nuclei.

We also reproduced the finding that higher-level, volatility PEs (equations 13 and 15) were represented in cholinergic regions. This time, however, we observed effects of volatility precision-weighted PEs in the cholinergic nuclei in the tegmentum of the brainstem, i.e., the pedunculopontine tegmental (PPT) and laterodorsal tegmental (LDT) nuclei (p<0.05 FWE voxel-level within an anatomical mask including all cholinergic nuclei) (Figure S5).

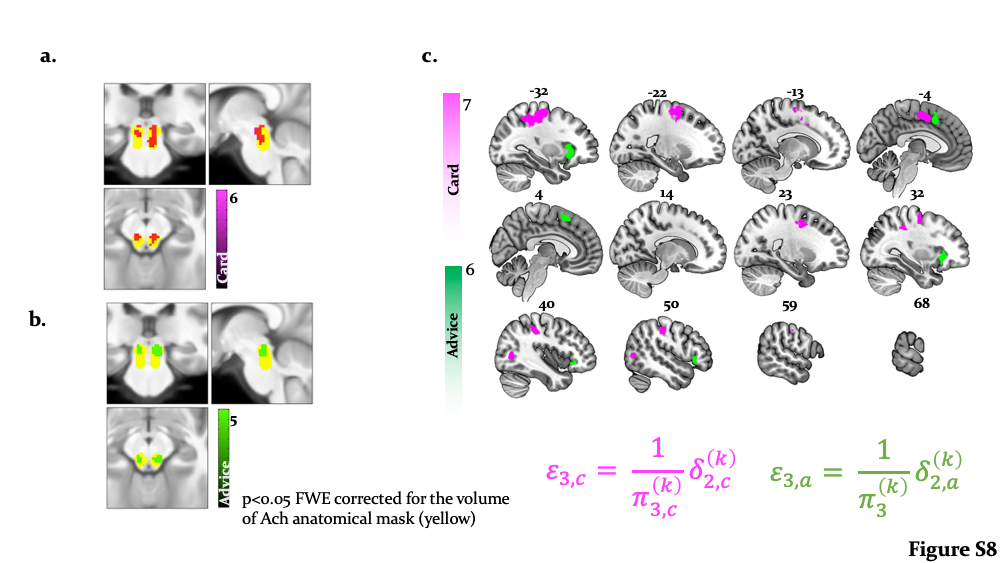

Figure S5| **Main effects of precision-weighted volatility**: Whole-brain activation by $\epsilon_{3}$ in the PPT/LDT nuclei: Activation of the bilateral cholinergic PPT/LDT associated with the signed precision-weighted volatility prediction error about the card (a) and the advisor fidelity (b) is shown at p<0.05 FWE corrected for the volume of our anatomical mask comprising cholinergic nuclei. Whole-brain activations by signed precision-weighted volatility PEs about the card colour probabilities were detected in the right inferior temporal gyrus, supramarginal gyrus, bilateral middle frontal cortex, and anterior cingulate sulcus. Whole-brain activations by signed precision-weighted volatility PEs about the advisor fidelity were detected in the right SMA, anterior insula and inferior lateral prefrontal cortex (whole-brain FWE cluster-level corrected, p<0.05).

### Supplementary Tables

|  | Hemisphere | x | y | z | *# Voxels* | *F-statistic* |
| --- | --- | --- | --- | --- | --- | --- |
| $\boldsymbol{\varepsilon}_{\boldsymbol{2}}$ card |  |  |  |  |  |  |
| lingual gyrus | R | 10 | -76 | -4 | 10994 | 157.31 |
| inferior occipital gyrus | L | -52 | -74 | 2 | 161 | 40.56 |
|  | R | 52 | -68 | 4 | 466 | 40.32 |
| insula | R | 32 | 22 | -4 | 934 | 53.36 |
|  | L | -34 | 22 | -2 | 745 | 27.62 |
| putamen | L | -30 | 0 | 4 | 331 | 21.44 |
| postcentral gyrus | L | -48 | -22 | 52 | 4230 | 82.43 |
| supramarginal gyrus | R | 58 | -20 | 42 | 1105 | 53.98 |
| dorsolateral PFC | R | 8 | 46 | 30 | 811 | 38.38 |
| substantia nigra | R | 8 | -18 | -14 | 3 | 14.09 |

**Table S1**: MNI coordinates and F-statistic of activations induced by precision-weighted prediction error about individually estimated card colour probability (equation 14). Related to Figure S3.

|  | Hemisphere | x | y | z | *# Voxels* | *F-statistic* |
| --- | --- | --- | --- | --- | --- | --- |
| $\boldsymbol{\varepsilon}_{\boldsymbol{2}}$ advice |  |  |  |  |  |  |
| cuneus | R | 12 | -88 | 20 | 6082 | 64.33 |
| insula | L | -32 | 20 | -8 | 620 | 42.37 |
|  | R | 32 | 22 | -4 | 663 | 42.18 |
| dorsal middle cingulate gyrus | L | -8 | -16 | 54 | 456 | 37.97 |
| putamen | L | -28 | -4 | 10 | 137 | 37.79 |
| dorsomedial PFC | L | -4 | 22 | 53 | 67 | 24.93 |
| posterior orbitofrontal cortex | L | -28 | 18 | -16 | 459 | 37.19 |
| dorsolateral PFC | R | 40 | 30 | 0 | 159 | 28.80 |
| ventral tegmental area | R | 4 | -16 | -12 | 151 | 49.99 |
|  |  | -4 | -18 | -14 | 150 | 37.94 |
| hippocampus | L | -20 | -28 | -4 | 55 | 33.69 |

**Table S2**: MNI coordinates and F-statistic of activations induced by precision-weighted prediction error about advice validity (equation 8). Related to Figure S4.
